# Bioengineering of a human innervated cardiac muscle model

**DOI:** 10.1101/2023.08.18.552653

**Authors:** Lennart Valentin Schneider, Bao Guobin, Aditi Methi, Ole Jensen, Kea Aline Schmoll, Michael Gani Setya, Sadman Sakib, Aminath Luveysa Fahud, Jürgen Brockmöller, André Fischer, Norman Y. Liaw, Wolfram-Hubertus Zimmermann, Maria-Patapia Zafeiriou

## Abstract

Cardiac autonomic neurons control cardiac contractility. Dysregulation of the autonomic nervous system can lead to sympathetic overdrive resulting in heart failure and an increased incidence of fatal arrhythmias. Here, we introduce innervated engineered human myocardium (iEHM), a novel model of neuro-cardiac junctions, constructed by fusion of a bioengineered neural organoid (BENO) patterned to autonomic nervous system and engineered human myocardium (EHM). Projections of sympathetic neurons into engineered human myocardium formed presynaptic terminals in close proximity to cardiomyocytes and an extensive vascular network co-developing in the tissues. Contractile responses to optogenetic stimulation of the accordingly engineered neuronal component demonstrated functionality of neuro-cardiac junctions in iEHM. This model will serve as a human surrogate system to delineate neuron and cardiac cell contribution to brain and heart diseases and is an important step towards engineering a human brain to heart axis in a dish.

## Introduction

Under physiologic conditions, sympathetic neurons (SN) arise from trunk neural crest ^1^ under the influence of morphogens such as retinoic acid, bone morphogenetic protein 4 (BMP4) and canonical Wnt activation ^1–3^. SN release noradrenaline, which binds to beta-adrenoreceptors present on cardiac cells (pacemaker or cardiomyocytes) to modulate contractility ^4^. In contrast parasympathetic neurons arising from cardiac neural crest release acetylcholine (ACh) and reduce beating rate ^4^.

Imbalance of autonomic neuronal regulation is a hallmark of brain and heart diseases (i.e. sudden cardiac death under epilepsy ^5^, catecholaminergic polymorphic ventricular tachycardia ^6^) as well as heart failure. Despite the importance of SN under physiologic and pathophysiologic conditions, their interaction with cardiomyocytes and other cells in the healthy and diseased heart is not well understood.

Human tissue engineering in combination with induced pluripotent stem cell (iPSC) ushered in a new era in human disease modeling. A number of tissue-engineered and organoid models of brain ^7–9^ or heart ^10–12^ have been developed and proved to efficiently model human diseases ^9,10,13^. A design principle in heart models is the co-culture of cardiomyocytes with non-myocytes, such as fibroblasts and endothelial cells ^10,14^. Similarly, in brain organoids neurons co-develop with glial cells ^7–9^. Co-culture experiments of cardiomyocytes and neurons have been performed with a demonstration of cell-cell interactions^2,3,15–17^. However, such models have not been advanced to the human tissue level, which has clear advantages in assessing cardiac muscle chronotropy or inotropy and deciphering neurocardiac interactions at the 3D syncytium level. Moreover, these models lack cellular complexity which may play a significant role in different pathomechanisms. For example, in the developing embryo sympathetic axons are chemoattracted towards the heart by nerve growth factor (NGF) released from perivascular cells and thus are located close to vessels ^18^. NGF is a key player in neuronal homeostasis in the heart ^19^ and its dysregulation after myocardial infarction is believed to underlie neuronal remodeling and sympathetic overdrive ^20^.

Aiming to generate a 3D neurocardiac interface, we patterned our previously established bioengineered neural organoid (BENO) ^7^ model to exhibit primarily SN activity. We then fused the resulting sympathetic neural organoid (SNO) with engineered human myocardium (EHM) ^10^ to allow SN to innervate and functionally interact with the cardiac cells.

## Results

### Generation of a sympathetic neural organoid (SNO)

SNO generation was achieved by patterning our previously established brain organoid model, BENO ^7^, towards sympathetic trunk fate ^2,3^. Embedded in a 3D collagen environment, human iPSC were committed to SN progenitors by modulation of pathways faithful to *in vivo* development. For the identification of an optimal protocol for derivation of SNO (**Figure 1A**) morphogen concentration, treatment window and cell number had to be fine-tuned. Optimized SNO comprised of 55±1% PHOX2B positive cells 15 days after initiation of directed differentiation (**Figure 1B, Supplementary Figure 1A**). When compared to BENO, SNO showed significantly greater levels of neural crest markers (*PHOX2B, ASCL1, GATA2*) and diminished expression of forebrain and hindbrain markers (*PAX6, GBX2* respectively, **Figure 1C, D, Supplementary Figure 1B**). By day 30, trunk neural crest cells differentiated into peripheral (PRPH) noradrenergic neurons expressing transcripts encoding for critical proteins for catecholamine synthesis and reuptake, such as dopamine beta hydroxylase and tyrosine hydroxylase (DBH, TH) as well as dopamine and noradrenaline transporters (*SLC18A2, SLC6A2*) (**Figure 1E-F, Supplementary Figure 2**). Both SNO and BENO contained similar levels of the pan-neural marker *TUBB3* (**Figure 1E**) confirming that the enhanced expression of noradrenergic markers in SNO reflected patterning to a sympathetic neural fate. Furthermore, transcript and immunofluorescence analyses revealed the presence of cholinergic neurons (*CHAT, SLC18A3*), glutamatergic neurons (*SLC17A6*), sensory neurons (*BRN3A*), and glia (*SLC1A3*), but absence of motor (*MNX1*) and GABAergic (*GAD1*) neurons (**Supplementary Figure 3**).

**Fig. 1.**
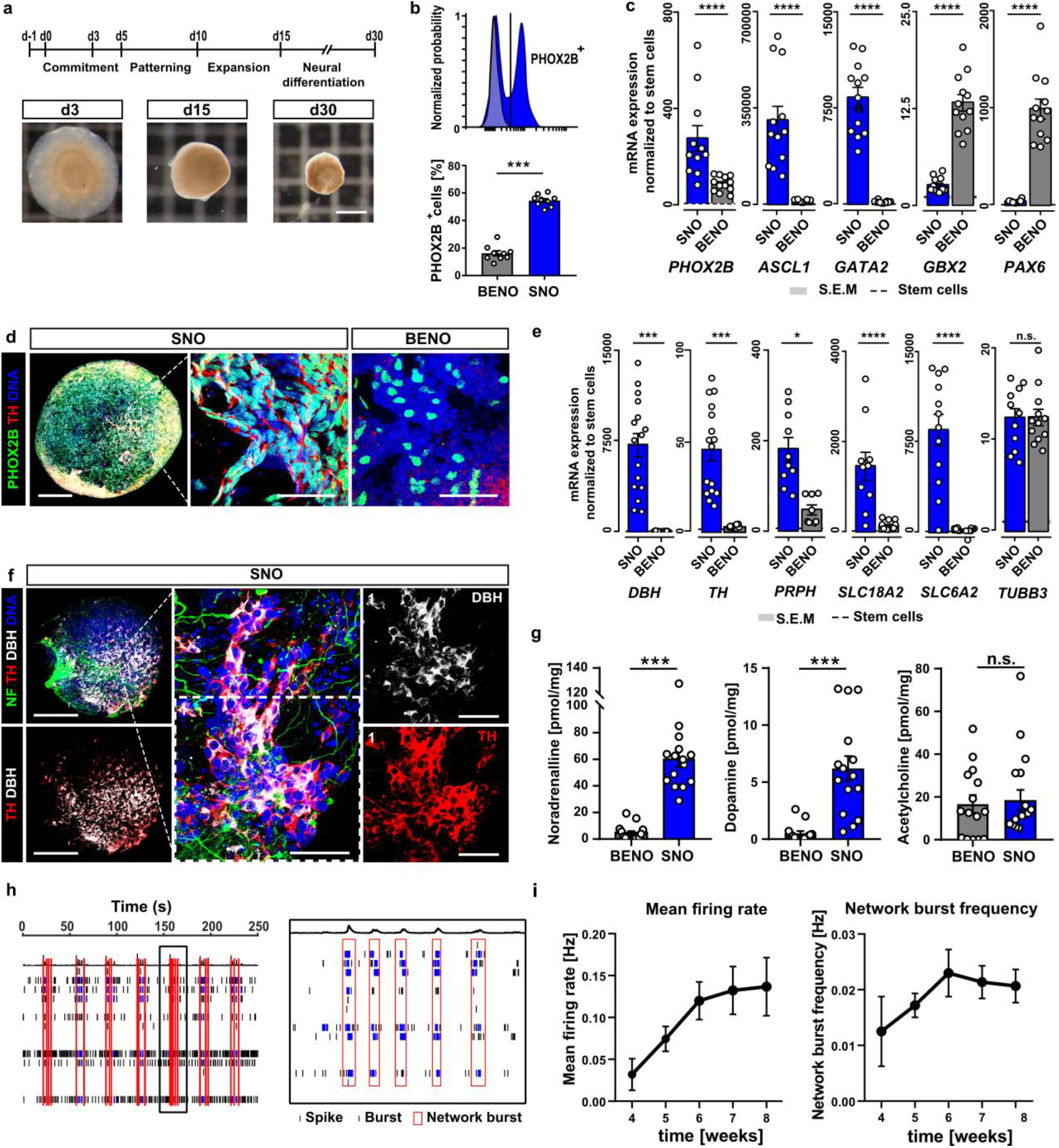
Generation and characterization of sympathetic neural organoids (SNO). **a**, **(Top)** Schematic of SNO differentiation protocol. **(bottom)** Brightfield images showing SNO at day (d) 3, 15 and 30 of development. Scale bar, 1 mm. **b**, Flow cytometry analysis of autonomic (PHOX2B^pos^) progenitor cells dissociated from SNO and BENO at d15. 3 independent differentiations, 3 biological replicates per differentiation (n=9). **c**, Expression analysis of autonomic progenitor marker (PHOX2B, ASCL1, GATA2) and cortical marker (GBX2, PAX6) for SNO and BENO (general brain organoid) at d15 normalized to GAPDH expression. SNO and BENO were normalized to expression in undifferentiated stem cells. 3 independent differentiations, 3-4 biological replicates per differentiation (n=11-12). **d**, Wholemount immunofluorescence analysis of d15 SNO and BENO stained for sympathetic progenitor marker PHOX2B (green) and TH (red). Scale bar, 500 μm. **(Right)** Close-up view of PHOX2B expression in SNO vs BENO. Scale bar, 50 μm. **e**, Expression analysis of pan-neural marker TUBB3, sympathetic markers (DBH, TH, SLC18A2, SLC6A2) and peripheral neuron marker peripherin (PRPH) for SNO and BENO at d41 normalized to GAPDH expression. SNO and BENO were normalized to expression in undifferentiated stem cells (dashed line). 3-4 independent differentiations, 3-4 biological replicates per differentiation (n=11-16), **f, (Left)** Wholemount immunofluorescence analysis of d41 SNO and BENO stained for sympathetic neuron marker DBH (grey), TH (red) and NF (green). Scale bar, 1 mm. **(Center)** Close-up view depicting sympathetic neurons co-expressing TH and DBH in SNO. Scale bar, 25 μm. **g**, LC-MS/MS analysis of BENO and SNO lysates at d41 normalized to wet weight of the tissues. 3 independent differentiations, 5 biological replicates per differentiation (n=15). **h, (Left)** Representative raster plot of SNO spontaneous activity at 8 weeks. **(Right)** Detail of a cluster of network bursts from the same recording (boxed in left panel). **i**, Timecourse of SNO mean firing rate and network burst frequency from weeks 4-8. n=46-48 SNO from 3 independent differentiations. All data is presented as mean ± s.e.m., Mann-Whitney test or unpaired, two-tailed Student’s t-test were performed, depending on normality of the data set.p*≤0.05, p***≤0.001, p****≤0.0001.

On day 40, liquid chromatography analysis of SNO vs BENO, showed significantly higher levels of sympathetic neurotransmitters noradrenaline (60.4±6.2 vs 4.4±1.5 pmol/mg) and its precursor dopamine (6.18±1.09 vs 0.51±0.20 pmol/mg), but no enrichment for acetylcholine (18.41±4.91 vs 16.73±4.19 pmol/mg), confirming a predominantly sympathetic nature of the neurons in SNO (**Figure 1G, Supplementary Figure 4**). SNO developed functional neuronal networks of increasing spontaneous activity (mean firing rate) and complexity (network burst frequency) that reached a plateau by 6 weeks of differentiation (**Figure 1H-I**) and demonstrated a concentration dependent response to nicotine (EC50 29.4±15.4 μM; **Supplementary Figure 5**). The latter is a hallmark for postganglionic neurons of the sympathetic nerve system and together with evidence for ACh synthesis, which is restricted to preganglionic neurons, suggests the formation of sympathetic ganglia in SNOs. Transcript and whole mount immunofluorescence analysis of SNO generated from three different iPSC lines indicated a highly reproducible protocol leading to similar levels of predominantly SN markers (**Supplementary Figure 6**).

### Generation and characterization of innervated human cardiac muscle (iEHM)

For the generation of the neurocardiac interface, 4-week old SNO were placed between the two arms of 4-week old EHM and the two tissues were allowed to fuse (**Figure 2A-C**). Innervated EHM (iEHM) exhibited spontaneous synchronous contractions from 2 days after fusion suggesting no inhibition of SNO to EHM function (**Supplementary video 1**).

**Fig. 2.**
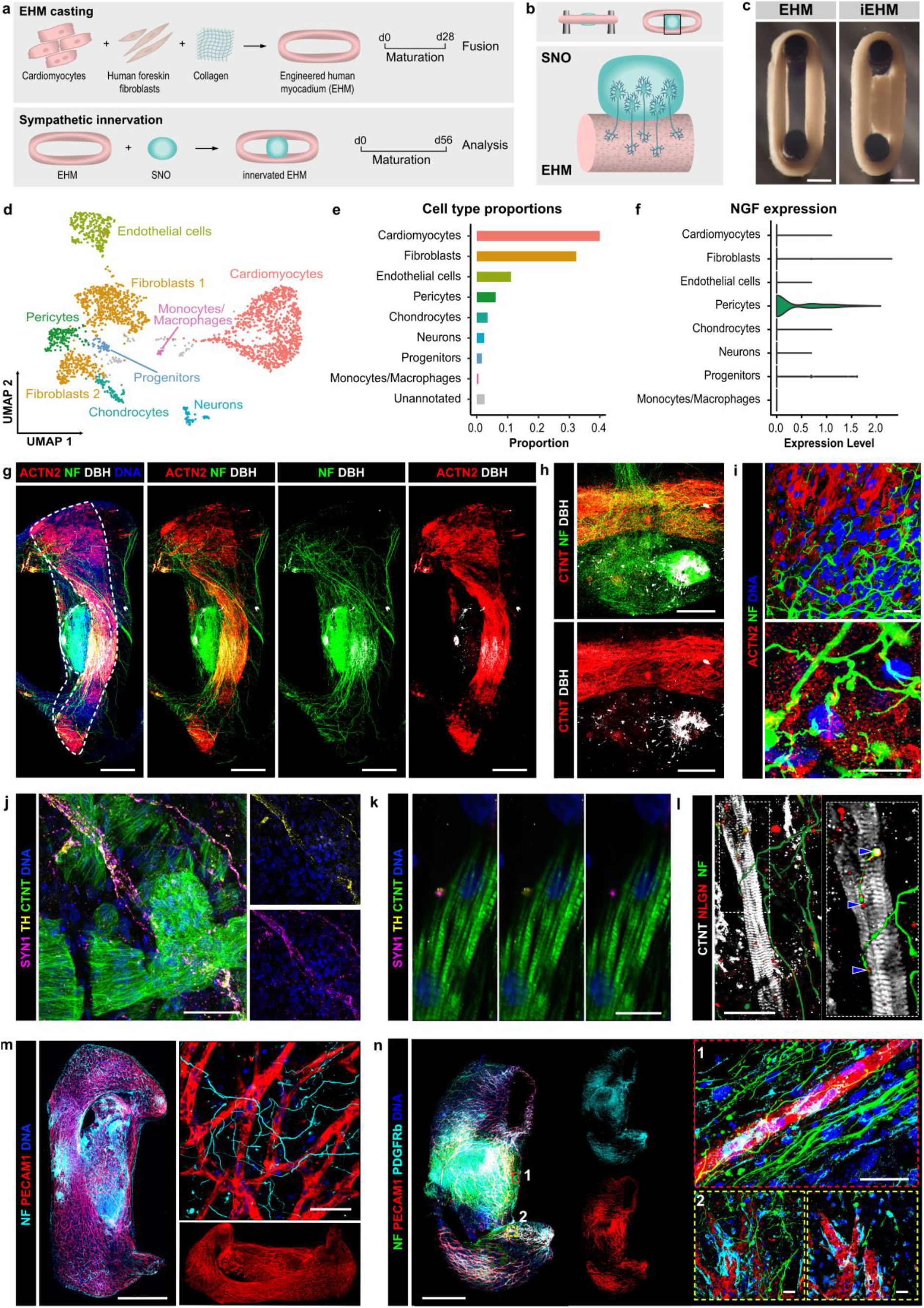
Innervated heart muscle (iEHM) generation and characterization. **a**, Schematic of iEHM generation workflow. **b**, Schematic of sympathetic innervation from the SNO projecting towards the EHM. **c**, Brightfield images of EHM and iEHM at 2 weeks of fusion. Scale bar, 1 mm. **d**, UMAP embedding and clustering of pooled snRNA-seq profiles from 8-week iEHM. Representative uniquely per cluster expressed genes are provided in **Supplementary Figure 8. e**, Proportions of the eight major cell types identified in iEHM by snRNA-seq analysis. **f**, Contribution of iEHM cell types to neural growth factor (NGF) expression. **g**, WhIF images of 8-week iEHM stained for neural marker NF (green), cardiomyocyte marker ACTN2 (red), SN-marker DBH (grey) and DNA (blue). Scale bar, 1 mm. **h**, Detail of the iEHM mid-section showing a cluster of DBH (grey) and NF (green)-positive SN in close proximity to the myocardium (CTNT, red), Scale bar, 500 μm. **i**, Close-up view of axons (NF, green) densely innervating cardiomyocytes (ACTN2, red).Scale bar, **(Top)** 20 μm, **(Bottom)** 10 μm. **j**, IF-staining of sympathetic (TH, yellow) presynaptic structures (SYN1, magenta) distributed throughout the myocardium (CTNT, green). Scale bar, 50 μm. **k**, Close-up view shows co-localization of SYN1 and TH on cardiomyocytes (green). Scale bar, 5 μm. **l, (Left)** IF-staining of iEHM for CTNT (grey), post-synaptic marker NLGN1 (red) and NF (green). Scalebar, 20 μm. **(Right)** Close-up view showing synaptic bouton-like varicosities co-stained with NLGN1 on the surface of cardiomyocytes, indicated by blue arrowheads. **m**, WhIF of 8 week iEHM stained for endothelial cell marker PECAM1 (red) and NF (cyan). Scale bar, 1 mm. **(Top right)** Close-up view of capillary structures interlaced by neuronal axons. Scale bar, 50 μm. **n**, WhIF of 8 week iEHM stained for DNA (blue), PECAM1 (red), NF (green) and PDGFRb (cyan). Scale bar, 1 mm. **ROI 1:** Close-up view illustrates the presence of PDGFRb-positive pericytes co-localizing with capillary tube-like structures as well as axons extending alongside. Scale bar, 20 μm. **ROI 2:** Representative image of pericytes (cyan) lining vascular structures (red), **(Left)** Maximum intensity projection, **(Right)** Single plane. Scale bar, 20 μm.

To define iEHM cell composition, we performed single nuclei RNA sequencing in one representative iEHM 6 weeks after fusion. Leiden clustering revealed 9 distinct populations (**Figure 2D, Supplementary Figure 8**). iEHM consisted of 40% cardiomyocytes, 32% fibroblasts, 11% endothelial cells, 6% pericytes, 4% chondrocytes, 3% neurons, 2% progenitors and 1% monocytes/ macrophages (**Figure 2E**). Cell identity was further substantiated by the detection of cell-type characteristic sodium, potassium and calcium channels (**Supplementary Figure 9**). Cardiomyocytes could be subclustered in ventricular-like (55%) and atrial-like cluster (22%) while 23% of the cells presented a mixed identity (**Supplementary Figure 10A-B**). Expression of *SHOX2, CACNA1D, GJA5* within the atrial cardiomyocytes suggested the presence of pacemaker-like cells. Moreover, one of the two fibroblast populations (*COL1A1, COL1A2, COL3A1* and *TBX18*) in iEHM stemmed from SNO-derived neural crest cells and was characterized by *PAX3, ZIC1* and *HOXB3* expression (**Supplementary Figure 10C-D**). Interestingly, similar to the native heart ^18^, pericytes were identified as the primary source in iEHM for nerve growth factor (*NGF*), a neurotrophin responsible for chemoattracting SN in the heart (**Figure 2F**).

6 weeks after fusion, immunofluorescence analysis showed an extensive axonal network marked by Neurofilament (NF) extending towards cardiac muscle expressing alpha-actinin; higher magnification images showed neurons interlacing cardiomyocytes (**Figure 2G-I, Supplementary Figure 11A, Supplementary video 2**). To visualize axonal synaptic boutons / varicosities in close proximity with cardiomyocytes we stained for the pre-synaptic marker Synapsin (SYN), while TH served as a marker for noradrenergic neurons. SYN^pos^-TH^pos^ noradrenergic varicosities were found in close proximity to cardiomyocytes (**Figure 2J-K, Supplementary video 3-7**), suggesting potential for neurocardiac communication. Furthermore, neuroligin 1, a postsynaptic marker expressed also by cardiomyocytes (**Supplementary Figure 11B-C**) was found on cardiomyocytes co-localized with varicosities as has been shown in the native heart ^22^ (**Figure 2I**).

Since superficial layers of the tissue displayed SYN^pos^-TH^pos^ noradrenergic axonal boutons that were not close to cardiomyocytes (**Supplementary video 8**) we reasoned that neurons may interact with other cell populations found in iEHM such as vascular cells. Indeed, immunofluorescence analysis of iEHM in week 6 after fusion, validated the presence of an extensive vascular network consisting of PECAM1 positive endothelial cells surrounded by PDGFRbeta positive pericytes (**Supplementary Figure 12A-B**), interlaced by neurons that seemed to follow the vessels recapitulating cardiac innervation during development (**Figure 2M-N, Supplementary videos 9-11**). Although superficial layers contained capillary like structures (diameter 5-10 μm) in deeper regions of the iEHM, vessels with diameter between 20-40 μm were apparent (**Supplementary Figure 12 C**). Note that the origin of the vascular network is both from the SNO and the EHM component; the latter deriving from cardiomyocyte-enriched directed differentiations from iPSC ^10^. Such populations comprise approx. 2% CD31-positive endothelial cells, which if not selected out by for example metabolic selection ^23^ can form capillary like-structures (**Supplementary Figure 12 B**) under standard EHM culture conditions^10^. Although SNO contained sparse endothelial cells, only one out of four tissues developed a vascular network when tissues were maintained in SNO media. Culturing SNO in the iEHM media, that contains vascular endothelial growth factor, increased the frequency of capillary network development as well as their amount (**Supplementary Figure 12 D**), suggesting that as expected iEHM vascularization is supported by VEGF.

### Autonomic neurons functionally connect with cardiomyocytes in iEHM

Since iEHM contained noradrenergic axonal bouton-like structures in close proximity to cardiomyocytes, we reasoned that the SN have an impact on tissue function and may form functionally relevant neuro-cardiac junctions. First, we investigated the contractile performance of iEHM vs EHM, 6 weeks after fusion. Isometric force measurements under electrical stimulation at 1.5 Hz showed no difference in iEHM and EHM contractile performance (**Figure 3A**), suggesting no effect of autonomic neurons in tissue contractility. However, when stimulated with inotropic agent isoprenaline, iEHM presented beta adrenoreceptor desensitization in comparison to EHM with significantly higher EC50 value (0.23±0.05 μM vs 0.10±0.03 μM respectively) (**Figure 3B**). Beta adrenoreceptors sensitivity to isoprenaline is inversely correlated with the presence of autonomic neurons ^24^ and noradrenaline *in vivo* ^25^, indicating that cardiomyocytes sense noradrenaline secreted in iEHM. Next, we investigated the effect of autonomic neurons on iEHM chronotropy by optogenetic means. We utilized a neuronal specific optogenetic iPSC line in which the red-shifted variant of channel rhodopsin f-Chrimson ^26^ was integrated in the AAVS1 locus under the human synapsin promoter (SYN_fChrimson). Opto-SNO showed higher firing rate and network burst frequency under light stimulation by week 8 of culture (**Figure 3C-D**). We next generated iEHM by fusing opto-SNO with wildtype EHM and compared them to standard EHM, to rule out unspecific activation of cardiomyocytes by light or heat (**Figure 3E**). Light stimulation of optogenetic iEHM evoked a positive chronotropic response up to two-fold of baseline (**Supplementary video 12**). Beating rate analysis of 42 iEHM (N=3) showed a mean beating rate increase of 24±4% in iEHM vs 1±3% in EHM, providing proof for functional connectivity between the autonomic neurons and pacemaker-like cells (**Figure 3F**). Pacemaker-like cells co-expressing *SHOX2/HCN4*, markers essential for pacemaker activity in the embryonic heart ^27^, were identified by snRNAseq (**Supplementary Figure 13A**) and immunofluorescence (**Supplementary Figure 13B**).

**Fig. 3.**
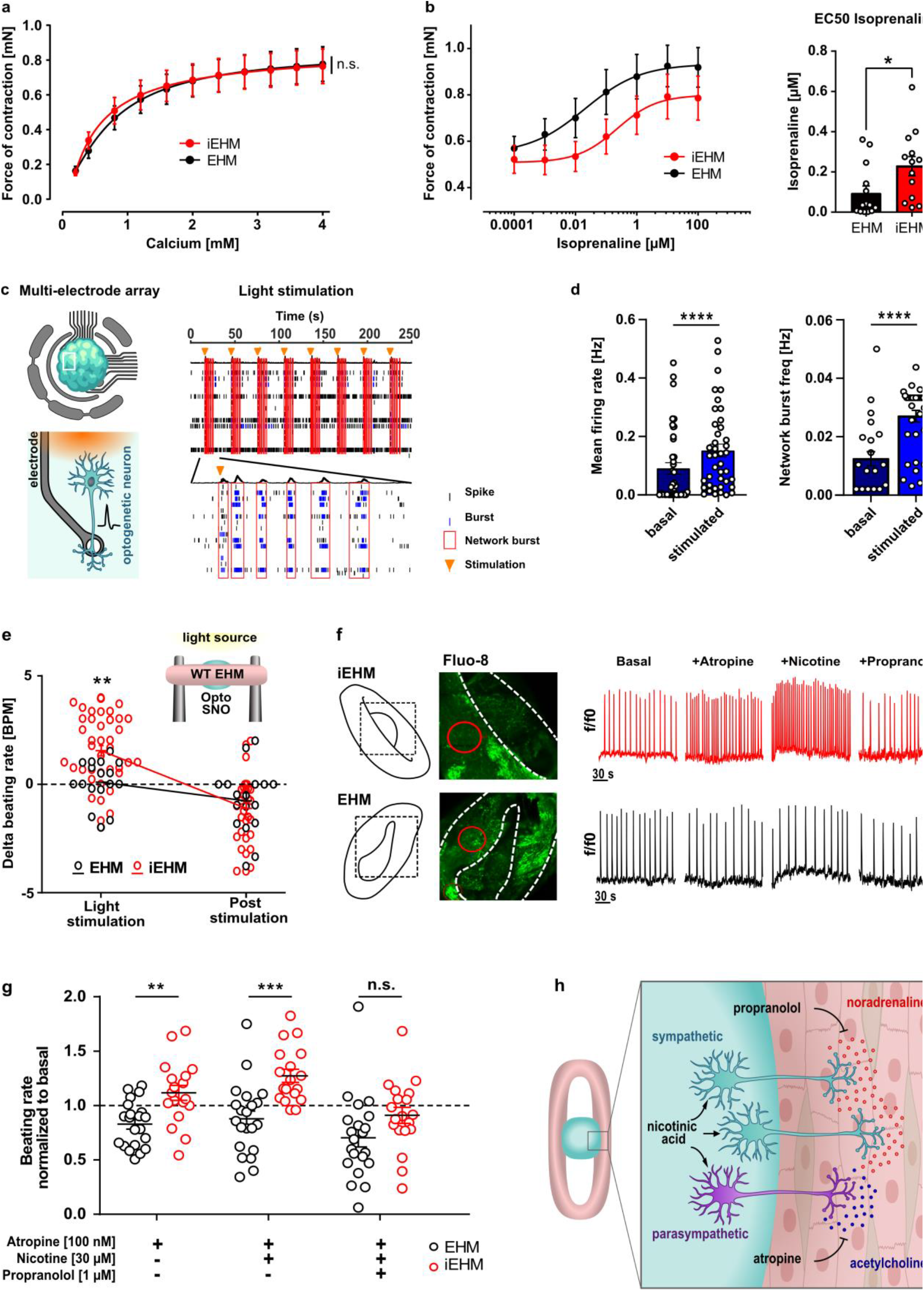
Functional analysis and connectivity between neurons and cardiac cells in iEHM. **a**, Force of contraction (FOC) recorded under increasing calcium concentrations and electric pacing at 1.5 Hz 57 days after fusion in EHM and iEHM. Three independent differentiations, 6-8 biological replicates per differentiation (n=22), ^ns^P=0.9278 by 2-way ANOVA. **b, (Left)** FOC response of EHM and iEHM to escalating concentrations of isoprenaline recorded at the EC50 of calcium and electrical stimulation at 1.5 Hz 57 days after fusion. **(Right)** Comparison of the EC50 of isoprenaline between EHM (n=13) and iEHM (n=14) from the same experiment. Two independent differentiations, 6-7 biological replicates per differentiation, *P=0.0125 by Mann-Whitney test. **c, (Left)** Schematic of optogenetic SNO mounted to multi electrode array (MEA), (below) schematic of red-light stimulation of optogenetic neuron in proximity to electrode. **(Right)** Example raster plot showing light-induced activity of SNO at 8 weeks of culture. Light stimulation pulses are indicated with orange triangles. **d, (Left)** Endpoint (8 weeks) measurement of basal and light-stimulated mean firing rate in optogenetic SNO. 3 independent differentiations, 12-15 biological replicates per differentiation (n=41), ****P<0.0001 by Wilcoxon test. **(Right)** Measurement of basal and light-stimulated network burst frequency in optogenetic SNO at the same date. n(stimulated)=36, n(unstimulated)=17, 3 independent differentiations, ****P<0.0001 by Mann-Whitney test. **e**, Schematic: Light stimulation of optogenetic SNO (opto SNO) leads to stimulation of the wildtype EHM in iEHM. Measurement of differences in beating rate of EHM and iEHM upon light stimulation. n(EHM)=15, n(iEHM)=39, 3 independent differentiations, age=5-10 weeks of fusion, **P=0.0022 by 2-way ANOVA with Sidak’s multiple comparisons post hoc test. **f, (left)** Representative confocal image of live recording of EHM and iEHM stained with calcium-sensitive dye Fluo-8 (green) including regions of interest (red). **(Right)** Traces of mean fluorescence intensity of Ca^++^-indicator dye Fluo-8 of iEHM and EHM upon pharmacological stimulation (more details in **Supplementary Figure 14B-C**). **g**, Quantification of EHM and iEHM beating rate during sequential treatment with 100 nM atropine, 30 μM nicotine and 1 μM propranolol. n(EHM)=21, n(iEHM)=19 (8 out of 27 iEHM were not responding, more details in **Supplementary Figure 14B**), 4 independent differentiations, age=8-10 weeks of fusion, Beating rate of the groups is normalized to the individual basal beating rate (indicated by dashed line). **P=0,0058 by Mann-Whitney test, ***P=0.0002, ^n.s.^P=0.0764 by unpaired t-test. Data are displayed as bar graphs with mean±s.e.m. as well as all individual data points. BPM=beats per minute, EHM=engineered human myocardium, f/f0=ratio of fluorescence intensity, iEHM=innervated EHM, SNO=sympathetic neural organoid. **h**, Schematic of pharmacological stimulation of iEHM: nicotinic acid stimulates neurotransmitter release from sympathetic and parasympathetic neurons in the iEHM. Effect of acetylcholine is inhibited by atropine administration and effect of noradrenaline is inhibited by non-specific beta adrenoreceptor blocker propranolol.

Furthermore, to delineate the contribution of the sympathetic and a potential parasympathetic component to iEHM beating response, we utilized classical pharmacological interventions (**Figure 3G**): (1) to block muscarinic receptors and thus ACh released by parasympathetic neurons (100 nM atropine), (2) to activate postganglionic nicotinergic receptors on sympathetic or parasympathetic neurons (30 μM nicotine) and (3) to block postsynaptic beta-adrenoreceptors (10 μM propranolol) expressed on pacemaker-like cells. The lack of a response to atropine in control EHM tissues showed that in the absence of autonomic neurons atropine does not directly affect cardiac muscle beating rate. In contrast, iEHM, showed a slight positive chronotropic response to atropine suggesting the blockade of parasympathetic neurocardiac junction (NCJ). Since nicotine would stimulate both sympathetic and parasympathetic neurons, we reasoned that pre-incubation with atropine would block the action of ACh and allow us to evaluate the absolute effect of SN activation only. Indeed, addition of nicotine in the presence of atropine, significantly increased beating rate by 38±13% of baseline (n=19, N=5). In the native tissue, postganglionic SN upon nicotine stimulation would release noradrenaline that would modulate chronotropy by binding to beta-adrenoreceptors in pacemaker-like cells. Blockade of beta-adrenoreceptors by propranolol abrogated the chronotropic effects of nicotine, providing further evidence for a functional NCJ in iEHM. 19 out of 27 iEHM responded to these stimulations, while 8 iEHM showed no response (**Supplementary Figure 14**), suggesting that catecholamine release and tissue level diffusion was insufficient to reach spontaneously beating cardiomyocytes or pacemaker-like cells in iEHM.

## Discussion

The brain communicates with the heart via the autonomic nervous system. Although increasing evidence suggests that channelopathies or miscommunication between brain and heart underlie a number of diseases ^5,6,28^ including heart failure, our understanding of human neuro-cardiac cell interaction and cardiac neuron plasticity is limited.

To study the functional interaction between autonomic neurons and cardiac cells, monolayer coculture systems ^2,3,17^ as well as microfabricated devices ^15,16^ have been employed. As a cell source, isolated primary cells were co-cultured with human iPSC-derived cells while only recently human derived co-cultures were developed ^15,17^. These systems allow cell-to-cell interaction, but provide limited information about developmental processes, cell organization in 3D as well as tissue dynamics. Three-dimensional cultures as organoids and engineered tissues have been shown to present higher cellular complexity, self-organization as well as enhanced maturation resembling the native tissue ^29,30^.

In this study, we bioengineered a human innervated cardiac muscle (iEHM) with functional neuro-cardiac junctions that emulates the cellular complexity of the native heart. Single nuclei sequencing showed that similar to adult heart data ^31^, iEHM consisted of cardiomyocytes, fibroblasts, endothelial cells, pericytes, neurons and monocytes. In addition, iEHM contained a small chondrocyte-like population that has not been reported in the adult human heart. Interestingly, neural crest-derived chondrocytes ^32^ have been reported in avian and mammalian embryonic heart ^33,34^ close to cardiac valves.

Resembling the epicardial innervation pattern of the native heart, iEHM innervation density showed an inverse correlation to the tissue depth. Interestingly, neuronal axons and presynaptic axonal boutons in iEHM were found in close proximity to capillary-like structures, which is in line with *in vivo* observations ^18^. The vascular network observed in iEHM, consisted of capillary-like (diameter 5-10 μm) and arteriole-like (diameter 20-40 μm) structures. The vascular cells originated from both the EHM component, which contained endothelial cells as a byproduct of mesoderm/cardiac differentiation and the SNO component. The development of endothelial cells in the SNO is not surprising since cKIT positive endothelial cells have been shown to emerge during cardiac neural crest development *in vivo* due to the transient BMP4 expression ^35^. Similarly, SNO are submitted to transient BMP4 exposure during the commitment phase. Another interesting point about the co-development of vessels and neurons is that during development, mural cells such as pericytes and smooth muscle cells produce and secrete NGF, which chemoattracts noradrenergic axons and guide sympathetic innervation of the heart ^18^. Similarly, in iEHM pericytes surrounding the dense capillary network were found to be the main source of NGF. Interestingly, NGF is considered one of the key factors orchestrating neuronal homeostasis in the adult heart. After myocardial infarction, NGF dysregulation is hypothesized to significantly contribute to neuronal remodeling, leading to sympathetic overdrive and arrhythmias ^36^.

As SN innervated the deeper layers of the EHM, TH positive synaptic varicosities were found in close proximity to cardiomyocytes or pacemaker-like cells, as previously observed in human postmortem heart samples and co-cultured cells ^22^.

Since morphological analysis suggested the presence of NCJ, we next investigated the functional connectivity between SN and cardiac cells. 6 weeks after fusion, iEHM demonstrated high contractile performance similar to EHM, suggesting that the presence of the neuronal component or media do not negatively impact cardiomyocyte function. Beta-adrenoreceptor desensitization to isoprenaline stimulation, typically observed *in vivo* in the presence of chronic NE treatment suggested that cardiomyocyte sense the NE released by sympathetic terminals.

From 6 weeks after fusion, optogenetic stimulation and pharmacological interventions resulted in a significant modulation of iEHM beating rate, demonstrating evidence for functionally relevant NCJ. Further, we observed responses to atropine, a known muscarinic receptor 2 blocker, suggesting the presence of cholinergic neurons. The lack of *MNX1* expression in SNO, a definite marker of spinal motor neurons, indicates that these cholinergic neurons cannot be motor neurons. One possibility is that these cells are post-ganglionic parasympathetic neurons that derive from cardiac neural crest, a region of the spine developing dorsally to the trunk neural crest from which SN arise. This data suggest that other relevant spine identities may co-develop during the caudalization of the progenitor cells which would also explain the co-development of sensory cells in SNO. Moreover, we cannot exclude that the cholinergic neurons we observe maybe pre-ganglionic SN, since no definite markers exist to our knowledge that would allow us to discriminate the two populations. Nevertheless, irrespective to their origin, both populations would release ACh that would stimulate muscarinic receptors in cardiac pacemakers and reduce tissue beating rate. Of note, 8 out of 27 iEHM did not respond, suggesting that the NCJ were underdeveloped. The primary reason for this observation may be that the pacemaker-like cell component in EHM is low (<10%) ^10^ or that SN axons may not in all cases be developed into sufficiently close proximity to mediate a change in the pacemaker I_f_ current (mediated via HCN4 channels).

In conclusion, iEHM is a novel neuro-cardiac interface that presents three major advantages to current state of the art. First, it allows the study of electrically excitable networks of human autonomic neurons, vascular cells, and cardiomyocytes in a 3D environment; second, it allows the quantification of chronotropic responses of cardiac muscle induced by optogenetic / chemical manipulations of neurons; finally, since tissues are generated separately and are fused into one, “mix&match” of wild type and mutant iPSC-derived tissues can be used in the future to delineate the cell type/organoid contribution to specific pathologies (SN, vascular cells or cardiomyocytes) and furthermore delineate how diseased cardiomyocytes affect healthy neurons and *vice versa*.

## Acknowledgements

The authors would like to especially thank Bastian Bues, Davide Ibatici, Petra Tucholla, Daria Reher and Iris Quentin for the excellent technical assistance. The authors would like to thank Tobias Moser and Thomas Mager for providing the pSYN-f-Chrimson plasmid and Laura Zelarayan for the AAVS1 locus donor plasmid. The authors would also like to especially thank Susanne Lutz for the interesting scientific discussions. This project was financially enabled by support of the German research foundation (DFG ZA 1217/3-1), by the German Center for Cardiovascular Research (DZHK) granted to M.P.Z. and by the Multiscale Bioimaging Cluster of Excellence (MBExC) granted to M.P.Z. and W.H.Z. W.H.Z. is supported by the DZHK (German Center for Cardiovascular Research), the German Federal Ministry for Science and Education (BMBF FKZ 161L0250A), the German Research Foundation (DFG SFB 1002 C04/S01, IRTG 1816, EXC 2067-1), and the Foundation Leducq (20CVD04). Generation of the GMP line LiPSC-GR1.1 (also referred to as TC1133 or RUCDRi002-A; lot number 50-001-21) was supported by the NIH Common Fund Regenerative Medicine Program, and reported in Stem Cell Reports.^35^ The NIH Common Fund and the National Center for Advancing Translational Sciences (NCATS) are joint stewards of the LiPSC-GR1.1 resource. A derivative from a GMP post-production working cell bank of the TC1133-line was made available to the Institute of Pharmacology and Toxicology at the University Medical Center by Repairon GmbH. The BIHi001-A-2 iPSC-line was kindly provided by Sebastian Diecke, Max Delbrück Center, Berlin.

## Author contributions

**L.V.S**. contributed to the design of the work; the acquisition and analysis of data; drafted and revised the manuscript. **G.B**. contributed to the design of the work; the acquisition, analysis, interpretation of functional data and revised the manuscript. **A.M**. Analyzed the snRNAseq data and drafted the associated method section. **O.J**. Established, performed and analyzed the neurotransmitter quantifications by LCMS. **K. A. S**. Generated and validated the optogenetic iPSC line (hSyn_Chrimson_ RUCDRi002-A). **S.S**. Performed the single nuclei isolation. **M.G.S**. contributed to the acquisition and analysis of quantitative PCR. **A.L.F**. performed WhIF stainings. **J. B**. contributed to the LCMS analysis and revised the manuscript **A.F**. contributed to the transcriptome analysis and revised the manuscript **N.L**. Contributed to the scientific know-how for engineered human myocardium technology, acquisition and analysis of data. **W.H.Z**. interpreted data and revised the manuscript. **M.P.Z**. contributed to the conception, the design of the work; the acquisition, analysis, interpretation of data; drafted and revised the manuscript.

**All authors** have approved the submitted version and agreed both to be personally accountable for the author’s own contributions and to ensure that questions related to the accuracy or integrity of any part of the work, even ones in which the author was not personally involved, are appropriately investigated, resolved, and the resolution documented in the literature.

## Materials and Methods

### Pluripotent stem cell culture

The hiPSC lines (1) RUCDRi002-A, (2) hSyn_Chrimson_RUCDRi002-A, (3) UMGi093-A, and (4) BIHi001-A-2 were maintained in StemMACS™ iPS-Brew XF (Brew, Miltenyi, #130-104-368) containing 100 U/mL penicillin and 100 μg/mL streptomycin (Thermo Fisher, #15140-122). iPSC were detached and singularised using EDTA solution (0.5 M EDTA/PBS) and replated on Matrigel (1:120, BD Bioscience, #354230) at a density of 8,000-12,000 cells/cm^2^, passaged twice weekly in the presence of ROCK inhibitor (Y-27632, 5 μM, Stemgent, #04-0012).

### Cardiomyocyte differentiation and purification

At a confluency of 80-90%, cells were submitted to cardiac differentiation as previously reported ^10^. d0-d3, cells were supplemented daily with Basal Serum Free Medium (BSFM: RPMI 1940 (Thermo Fisher, #61870-044), 2% B27 Supplement (Thermo Fisher, #17504-044), 200 μM L-ascorbic acid-2-phosphate sesquimagnesium salt hydrate and 100 U/mL penicillin, 100 μg/mL streptomycin (Thermo Fisher, #15070063)) containing 9 ng/mL Activin A (R&D systems, #338-AC), 5 ng/mL BMP4 (R&D systems, #314-BP), 1 μM CHIR99021 (Stemgent, #04-0004) and 5 ng/mL bFGF (Miltenyi Biotech, #130-093-841). D3-d13, the cells were cultured in BSFM supplemented with 5 μM IWP4 (Stemgent, #04-0036) and d13-d17, cells were cultured in unsupplemented BSFM, which was changed every other day. For metabolic selection, cardiomyocytes were cultured in RPMI without glucose and glutamine (Thermo Fisher, #11879020) containing 2.2 mM sodium lactate (Sigma, #71723), 100 μM β-mercaptoethanol (Invitrogen, #31350-010), 100 U/mL penicillin, and 100 μg/mL streptomycin (Thermo Fisher, #15070063)) for 5 days.

### EHM generation

EHM were generated as previously described ^10^.

### BENO and SNO generation

BENO were generated as previously described ^7^. SNO differentiation was based on the BENO protocol and was optimized by testing concentrations and treatment windows for various morphogens.

### Whole-mount immunofluorescence staining

Tissues were fixed with 4% formaldehyde solution (Histofix, Carl Roth) for 2 h at 4 °C. Subsequently, they were washed twice with PBS and blocked for 30 min at 4 °C with staining buffer (StB; 5% FBS, 1% BSA, 0.5% Triton X-100 in PBS). Tissues were incubated with primary antibodies diluted in StB for 2 days at 4 °C (100 μl/tissue). Upon washing with StB for 6–8 h, tissues were incubated with secondary antibodies and Hoechst 33342 (Sigma) for another 2 days at 4 °C. After StB washings for a total of 6–8 h BENOs were mounted on glass coverslips with Moviol mounting medium (10% Fluoromount Mowiol 4-88 (Carlroth, #0713.1), 25% Glycerol (Merck-Millipore, #56-81-5), 0.1 M Tris (ITW Reagents, #A1086), 0.2% n-propyl gallate (Sigma-Aldrich, #P3130)). WmIF was visualized using confocal imaging performed on a Zeiss LSM 710 confocal microscope equipped with ZEN 2010 software and further analysed by ZEN and ImageJ. An antibody list with respective dilutions is provided in Supplementary Table 3.

### Tissue digestion and flow cytometry

For SNO and BENO digestion, d15 organoids were sequentially incubated for one hour with collagenase solution (PBS +Ca/Mg, 0.25% Collagenase Type I (Sigma-Aldrich, #C0130), 20% FBS (Thermo Fisher, #A4766801)) and Accutase dissociation reagent (97% StemPro® Accutase® Cell Dissociation Reagent (Merck-Millipore, #SCR005), 0.025% Trypsin (Thermo Fisher, #15090-046), 20 μg/mL DNaseI (Calbiochem, #260913). The digested cells were fixed in 4% formaldehyde at room temperature (RT) for 15 min and subsequently labelled with anti-PHOX2B primary antibody (Santa Cruz, #sc-376997, 1:250) and Hoechst 33342 in StB for 45 minutes at 4 °C. To remove the primary antibody, the cells were centrifuged at 3,000 xg, supernatant was discarded and the cells were resuspended and incubated in StB for five minutes at 4 °C. This step was repeated three times. Subsequently, cells were incubated with secondary antibody Alexa Fluor 488 (Invitrogen, #A32723, 1:1000) for 30 minutes at 4 °C under the exclusion of light. Following another 3x 5 minutes washing step, the cells were analysed using a BD LSR II flow cytometer (BD Biosciences) and FACSDiva (BD Biosciences) as well as FLOWJO software.

### Liquid chromatography-coupled tandem mass spectrometry (LC-MS/MS)

SNO were lysed in 500 μl precipitating agent (80% acetonitrile and 20% water) including noradrealine-d6, choline-d9 and buformin as internal standard for noradrenaline, acetycholine and dopamine respectively. Following 15 minute incubation at RT, 400 μL of the liquid phase were evaporated under a nitrogen stream. The residues were reconstituted in 400 μl of 0.1% (v/v) formic acid and 10 μl were injected into the LC-MS/MS system. Noradrenaline, dopamine and acetylcholine (ACh) content of SNO were analyzed by high-performance liquid chromatography (HPLC)−MS/MS using a Shimadzu Nexera HPLC system with a LC-30AD pump, a SIL-30AC autosampler, a CTO-20AC column oven, and a CBM-20A controller (Shimadzu, Kyoto, Japan). Separation was achieved on a Imtakt-Intrada Amino Acids Separation Column, 100 x 3.0 mm (ChromTech, #WAA34) with a C-18 guard pre-column. The HPLC was run at a low speed of 600 μL/min with an oven temperature of 37 °C. The mobile phase consisted of 29.85% 200 mM ammonium formate, 0.15% (v/v) formic acid and 70% (v/v) methanol. The HPLC system was coupled to an API 4000 tandem mass spectrometer (SCIEX, Darmstadt, Germany). Buformin (internal standard), Noradrenaline-d6 (internal standard), choline-d9 (internal standard) were quantified using the detection parameters given (Supplementary Table 1). Analysis was performed using the Analyst software (version 1.6.2, SCIEX, Darmstadt, Germany) and standard curves obtained by weighted linear regression. Representative spectra and standard curves are provided in **Supplementary Figure 4**.

### Quantitative real-time (qRT) PCR

Total RNA was extracted using the NucleoSpin RNA Mini kit (Macherey Nagel, #740955). 250 ng to 1 μg of RNA were reverse transcribed using M-MLV Reverse transcrtiptase (Promega, #M1705) and oligo(dT) primer. qRT-PCR was performed using Takyon Blue dTTP Master Mix (Eurogentec, #UF-NSMT-B0701). The primer sequencesare provided in Supplementary Table 2. Analysis was performed by 7900HT Real Time PCR System (Applied Biosystems) and the Sequence Detection System (SDS) v2.4 software (Applied Biosystems).

### Calcium imaging

The iEHM were stained with 1 μg/ml Fluo-8-AM (ABCAM, #ab142773) in carbogenated artificial cerebrospinal fluid (ACSF) buffer (26 mM NaHCO_3_, 10 mM Glucose, 1 mM MgSO_4_ 7H_2_O, 1.25 mM NaH_2_PO_4_, 2.5 mM KCl, 126 mM NaCl, 2 mM CaCl_2_) at pH7.3 for 15–30 min. During calcium imaging, iEHM were continuously perfused with ACSF at 37 °C. Fluorescence was recorded at 2-5 Hz and measurements were performed using Zeiss LSM 780 confocal microscope and ZEN 2010 software. The stimulation of ACh to muscarinic receptors was blocked by addition of 10 nM Atropine (Sigma Aldrich). In the presence of Atropine, 30 μM Nicotinic acid was used to stimulate nicotinic receptors on autonomic neurons. Beta-adrenoreceptor blockade was achieved by adding 10 μM Propranolol. Analysis was performed using Matlab R2012a (The Math Works, USA).

### Force of contraction analysis

Isometric force measurements of EHM / iEHM were perfomed in an organ bath setup as described previously ^10^. EHM and iEHM were mounted on two opposing hooks and submerged in normal Tyrode’s solution (120 mM NaCl, 5.36 mM KCl, 0.2 mM CaCl_2_, 10.5 mM MgCl_2_, 22.61 mM NaHCO_3_, 4.2 mM NaH_2_PO_4_, 5.55 mM Glucose, 0.57 mM L-Ascorbic Acid) at 37°C with a continuous flow of carbogen from beneath. Force changes were measured by a linear isometric force transducer mounted to the upper hook. The basal tension of the tissues was adjusted to 0.1 mN using a micromanipulator. On either side, a field electrode was placed for electrical pacing of the tissues.

### Electrical field potential measurements by multielectrode-array (MEA)

SNO were embedded in 3% low melting agarose and sectioned using Leica VT1000S vibratome (Leica) into slices of 400 μm diameter. SNO slices were mounted on Lumos MEA 48-well plates (Axion BioSystems, #M768-tMEA-480PT) coated with 1:30 Matrigel. Gold grids for transmission electron microscopy (Gilder, #G50HEX-G3) were used to stabilize the tissue. The measurements of electrical field potentials were performed by Maestro Pro (Axion BioSystems) at 37 °C and 5% CO2. The Lumos system (Axion BioSystems) enabled optogenetic stimulation of the organoids by delivering a pentaphasic light pulse with a central wavelength of 612 nm and an intensity of 1.54 mW/mm^2^ every two minutes for one hour. Stimulation experiments were performed by addition of increasing concentrations (1 μM-1 mM) of nicotine diluted in pre-warmed Basal medium, while the added volume never exceeded five percent of the wells’ media volume. Analysis was performed by the Neural Metrics and Plotting Tool software (Axion BioSystems).

### Single nuclei RNA sequencing

Frozen iEHMs were used to isolate nuclei, adapted from Sakib *et al*.^37^ with modifications. In brief, 500 μL EZ prep nuclei lysis buffer (Sigma, #NUC101) were added to the frozen samples and dounce homogenized for 40-60 strokes (until not big tissue chunks were seen) using plastic pestles in a 1.5 mL DNA low bind tube. Lysates were transferred into a 2 mL DNA low bind tube and additional lysis buffer were supplemented to fill up to 2 mL mark. Samples were incubated for 5 minutes on ice and later centrifuged for 500 xg for 5 minutes at 4 °C. Supernatants were discarded and the nuclei pellets were resuspended into 2 mL lysis buffer, and incubated on ice for 5 minutes. After centrifugation (500 xg, 5 minutes, 4 °C), the pellet was resuspended into Nuclei Suspension buffer, NSB (0.5% OmniPur BSA (Merck Millipore, #2905), 1:200 RNasin® Ribonuclease Inhibitor (Promega, #N2615), 1x cOmplete™, EDTA-free Protease Inhibitor Cocktail (Roche, #04693132001), dissolved in 1x PBS (RNAse free, Invitrogen, #AM9624). Filtered through 0.22 μm Syringe Filter Unit (Millipore, #SLGVR33RB)). It was centrifuged again for 500 xg, 5 minutes at 4 °C, and resuspended into 500 μL NSB. The lysate was strained through 40 μM Flowmi® Cell Strainers (Sigma, #BAH136800040) into a FACS tube. 1:500 of 7-AAD (Invitrogen, #00-6993-50) nuclei staining solution was added and nuclei were gated from the debris (**Supplementary Figure 7**) and sorted with a BD FACSaria III with 85 μM Nozzle, in a 15 mL falcon tube, coated with and containing 1 mL NSB. Around 200,000 nuclei were sorted per sample. Sorted iEHM nuclei were immediately taken to process for single nuclei barcoding using 10X Genomics Chromium Single Cell 3’ Reagent Kits (v3 Chemistry). 5000 nuclei were subjected to GEM generation and barcoding using Chromium Controller, according to the manufacturer’s protocol. For cDNA amplification, 14 PCR cycles were used. Indexed cDNA libraries were pooled into a single pool and sequenced 4 times in Illumina NextSeq 550 to achieve >50,000 reads/nuclei. Raw BCL files were demultiplexed, mapped and counts files were generated using cellranger-4.0.0, using hg38 pre-mRNA reference genome.

### Single nuclei RNA-sequencing data analysis

Cell Ranger (10x Genomics) was used for demultiplexing, alignment and generation of gene counts. The dataset was filtered to remove nuclei where less than 200 genes were detected, as well as genes that were expressed in less than 10 nuclei. Any nuclei expressing more than 1% mitochondrial gene counts were also removed. The data were normalized using the sctransform ^38^ package (version 0.3.2) from the Seurat ^39^ (version 4.0.2) toolkit in R (version 4.0.1). DoubletFinder ^40^ (version 2.0.3) was used to detect and remove any potential doublets. Leiden clustering ^41^ was performed using the FindClusters() function in Seurat and Uniform Manifold Approximation and Projection (UMAP) ^42^ was used for visualization of clusters. The top markers for each cluster were detected using the FindAllMarkers() function in Seurat. Cell-type annotation of clusters was done using the expression of selected marker genes as listed in **Supplementary Figure 8**. The same analysis pipeline was followed for separate analyses of specific cell clusters after subsetting the original dataset accordingly.

## Statistical analysis

Statistical testing was performed in GraphPad Prism 9. All values are displayed as mean ± s.e.m Statistical tests were chosen depending on parametricity of the sample distribution, which was determined by Shapiro-Wilk and F-test. The respective tests as well as sample size (n) and number of independent experiments (N) are mentioned in the figure legends. Statistically significant differences were assumed with a probability value of *P*<0.05. Exact *P*-values are given in the figure legends.

