## Supplementary data for "Bioengineering of a human innervated cardiac muscle model"

### Table of content

#### Supplementary Figure


- **Supplementary Figure 1|** Sympathetic neural progenitor analysis in d15 SNO
- **Supplementary Figure 2|** Sympathetic neuron immunofluorescence analysis in d30 SNO
- **Supplementary Figure 3|** Identification of non-SN cells co-developing in SNO
- **Supplementary Figure 4|** Neurotransmitter quantification by LC-MS/MS analysis
- **Supplementary Figure 5|** Effect of nicotinic acid on electrical activity of SNO d73-87
- **Supplementary Figure 6|** Reproducibility of SNO differentiation protocol in three iPSC-lines
- **Supplementary Figure 7|** Gating strategy for sorting nuclei from frozen iEHM lysates using 7AAD staining
- **Supplementary Figure 8|** Cell cluster annotation of snRNA-sequencing
- **Supplementary Figure 9|** Expression analysis of ion channels across the cell types from snRNA-sequencing
- **Supplementary Figure 10|** Analysis of cardiomyocyte and fibroblast subcluster in the iEHM
- **Supplementary Figure 11|** Investigation of neuro-cardiac junctions in iEHM
- **Supplementary Figure 12|** Vascular network development in SNO, EHM and iEHM
- **Supplementary Figure 13|** Identification of pacemaker-like cells in iEHM
- **Supplementary Figure 14|** Details of pharmacological stimulation of iEHM


#### Supplementary Materials


- **Supplementary Table 1|** Detection parameters for a second compound-specific mass transition used as qualifier
- **Supplementary Table 2|** List of primer for quantitative real-time PCR
- **Supplementary Table 3|** List of primary and secondary antibodies


#### List of Supplementary Videos

### Supplementary Figures

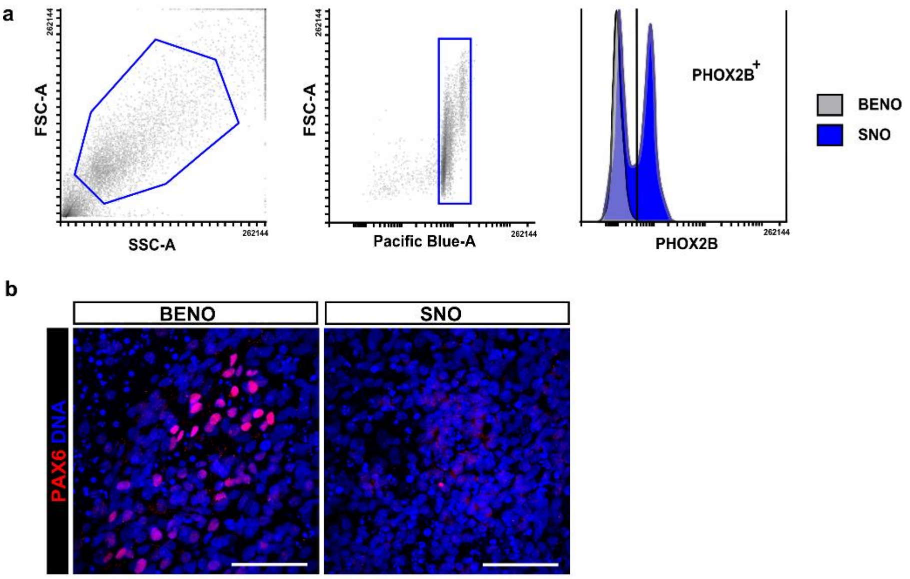


***Supplementary Figure 1| Sympathetic neural progenitor analysis in d15 SNO. a**, Gating of flow cytometry analysis of d15 neural progenitor cells isolated from SNO and BENO stained for DNA and PHOX2B. SSC-A=side scatter-area, FSC-A=forward scatter-area.* ***b,*** *Immunofluorescence of d15 SNO and BENO stained by cortical marker PAX6 (red). Scale bar, 50 µm.*

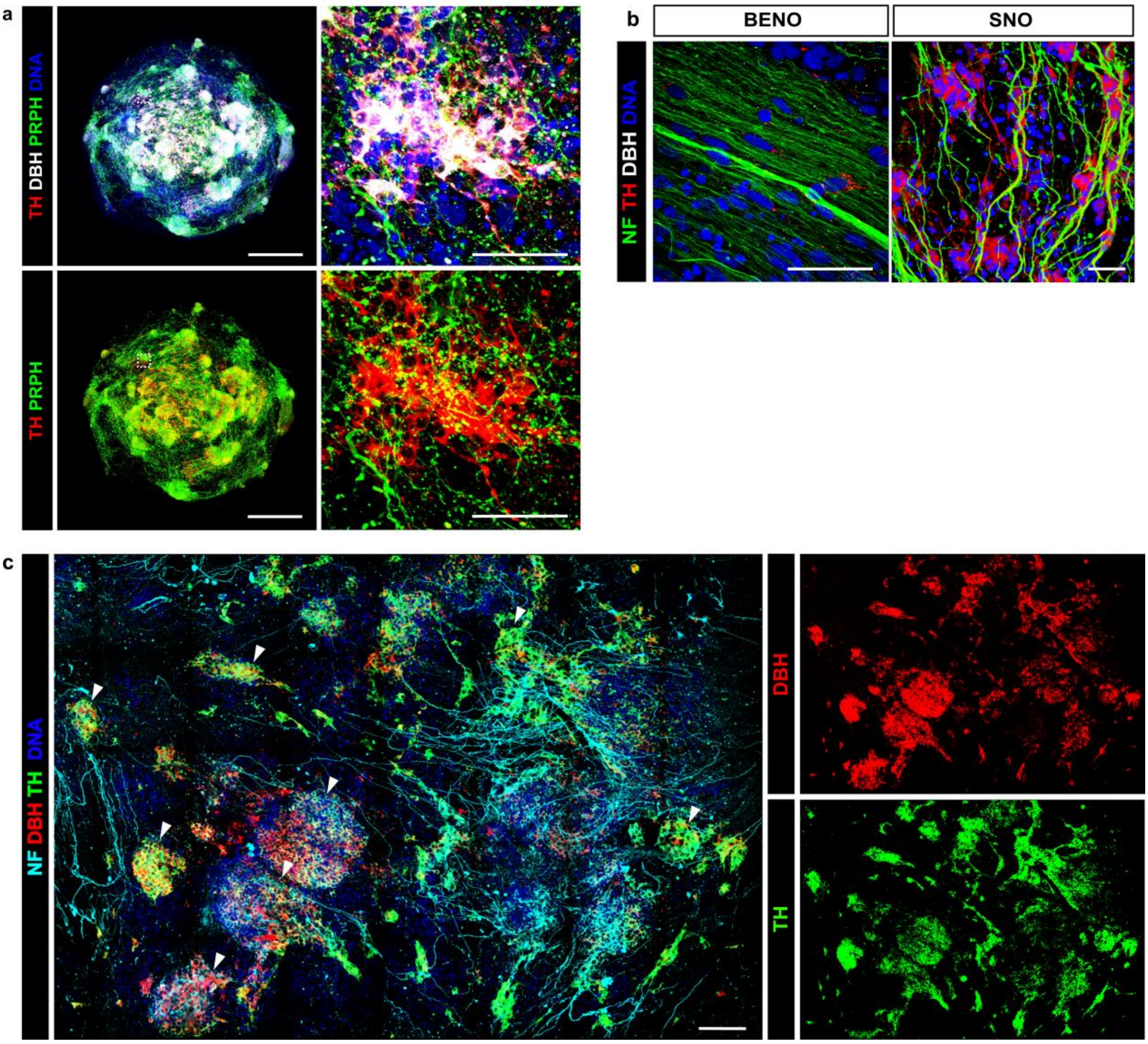


***Supplementary Figure 2| Sympathetic neuron immunofluorescence analysis in d30 SNO. a,*** *Representative image of d41 SNO stained for sympathetic marker TH (red), DBH (greys), peripheral neuron marker PRPH (green), and DNA (blue). Scale bar, 500 µm.* ***(Right)*** *Close-up view. Scale bar, 50 µm.* ***b****, Close-up view shows sympathetic neurons co-stained by DBH (red), NF (green) and DNA (blue) in SNO showing ganglia-like structures* ***(Right)****. BENO did not contain sympathetic neurons* ***(Left)****. Scale bars, 50 µm.* ***c****, High resolution imaging of SNO stained for sympathetic marker TH (red) and DBH (grey) as well as NF (green) and DNA (blue) showing widespread sympathetic ganglia-like structures (marked by white arrowheads). Scale bar, 100 µm.*

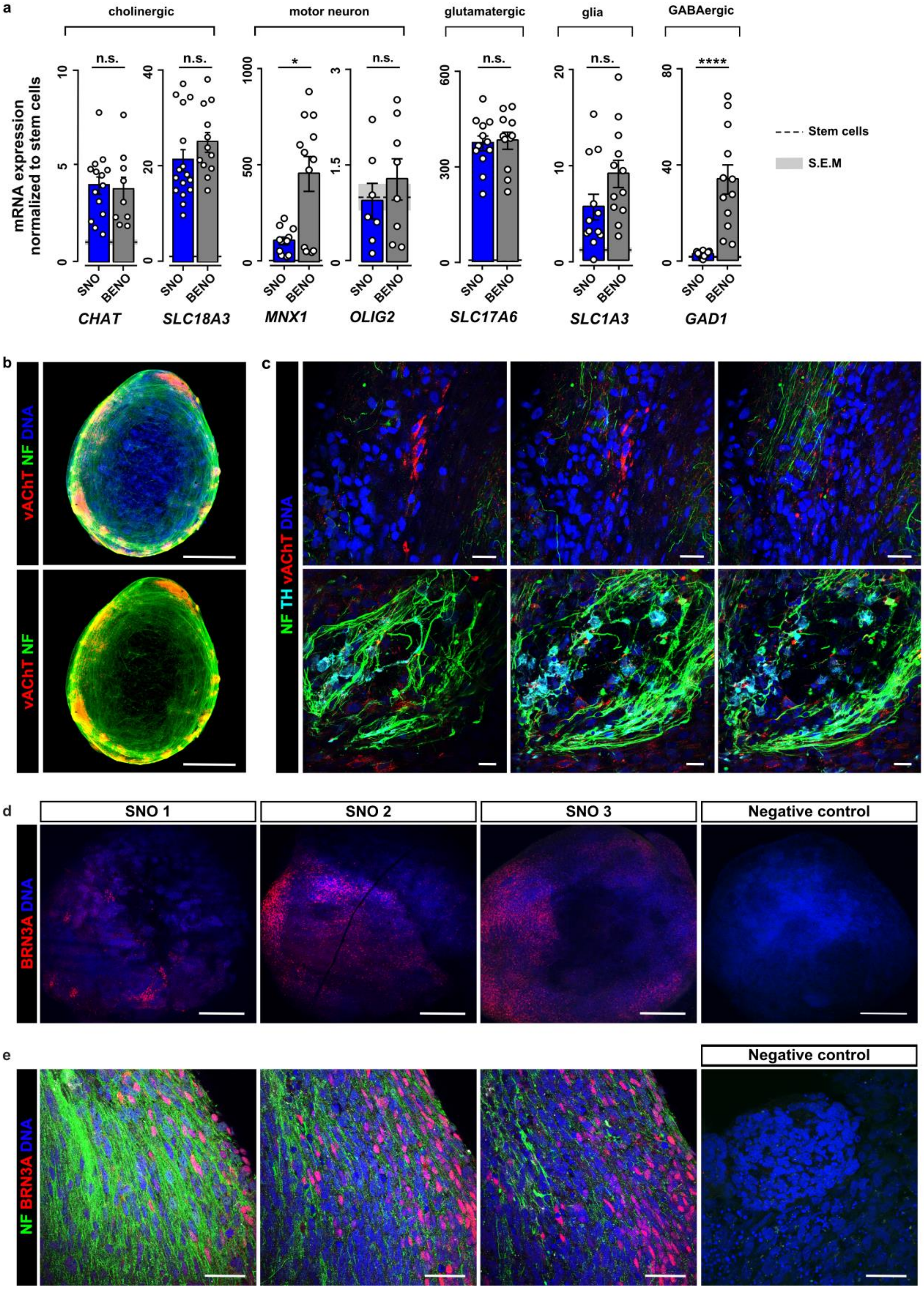


***Supplementary Figure 3| Identification of non-SN cells co-developing in SNO. a,*** *Expression analysis of cholinergic marker (CHAT, SLC18A3 [aka VACHT]), GABAergic (GAD1), motor (MNX1, OLIG2), glutamatergic (SLC17A6 [aka VGLUT2]) neuron marker and astrocyte (SLC1A3 [aka GLAST]) marker for SNO and BENO at d41 normalized to GAPDH expression. SNO and BENO were normalized to expression in undifferentiated stem cells. Mann-Whitney test or Student's t-test were performed, depending on normality*

*of the data set. 3-4 independent differentiations, 3-4 biological replicates per differentiation (n=9-16) p*≤0.05, p***≤0.001, p****≤0.0001.* ***b****, WhIF of d30 SNO stained for cholinergic marker vAChT (red), NF (green) and DNA (blue). Scale bar, 500 µm.* ***c,*** *Close-up view of cholinergic neurons vAChT (red) and sympathetic neurons marked by TH (cyan) from two different SNO regions. Scale bar, 20 µm.* ***d,*** *IF-staining of three individual SNO at d41 for sensory neuron marker BRN3A (red) and nuclear counterstaining (blue).* ***(Far right)*** *d41 SNO as no primary antibody control. Scale bar, 200 µm.* ***e,*** *IF of three planes from a representative region of d41 SNO stained for sensory neurons (BRN3A (red) and NF (green)). No primary antibody control can be seen on the right. Scale bar, 20 µm.*

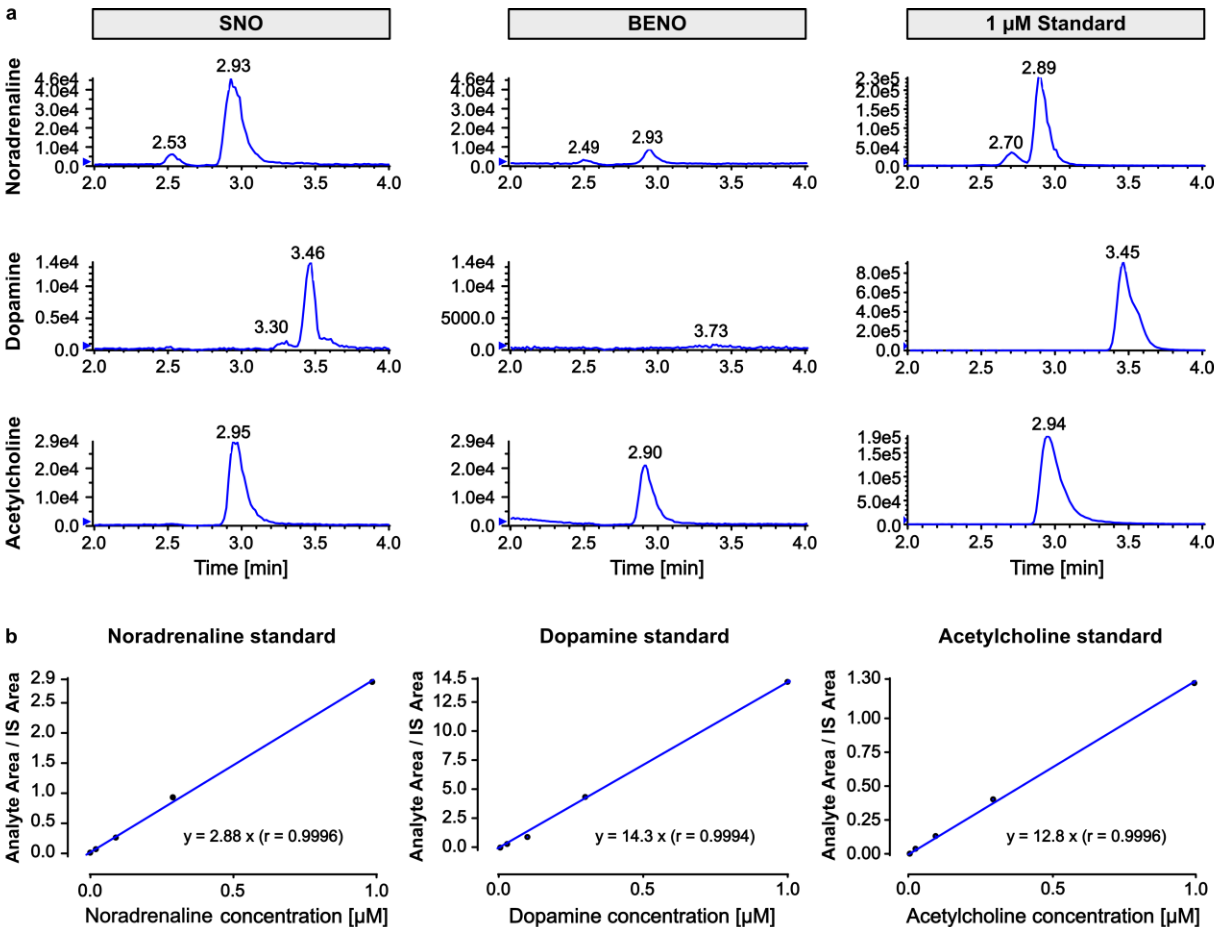


***Supplementary Figure 4| Neurotransmitter quantification by LC-MS/MS analysis. a,*** *Representative LC-MS/MS spectra from lysed SNO and BENO for the noradrenaline, dopamine and acetylcholine compared to the respective standard at 1 µM concentration.* ***b,*** *Standard curves of all three neurotransmitters (concentrations from 1 µM to 0.001 µM). Linear regression ("through zero") was calculated for the integrated peaks and used for quantification of the sample concentrations. Y-intercept and correlation coefficient r are indicated in the graphs.*

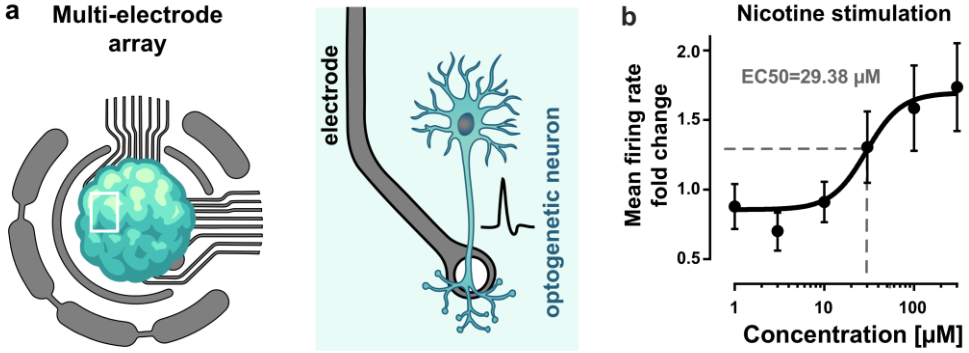


***Supplementary Figure 5| Effect of nicotinic acid on electrical activity of SNO d73-87. a***, *Schematic of the placement of SNO on the multi-electrode array.* ***b***, *Dose response curve of mean firing rate of SNO upon stimulation with escalating concentrations of nicotine normalized to basal firing rate and untreated control. $EC_{Nicotine}50$=29.38 µM, Nonlinear fit using variable slope (four parameters, ordinary fit). 3 independent differentiations, 6-7 biological replicates per differentiation (n=19), mean ± s.e.m.*

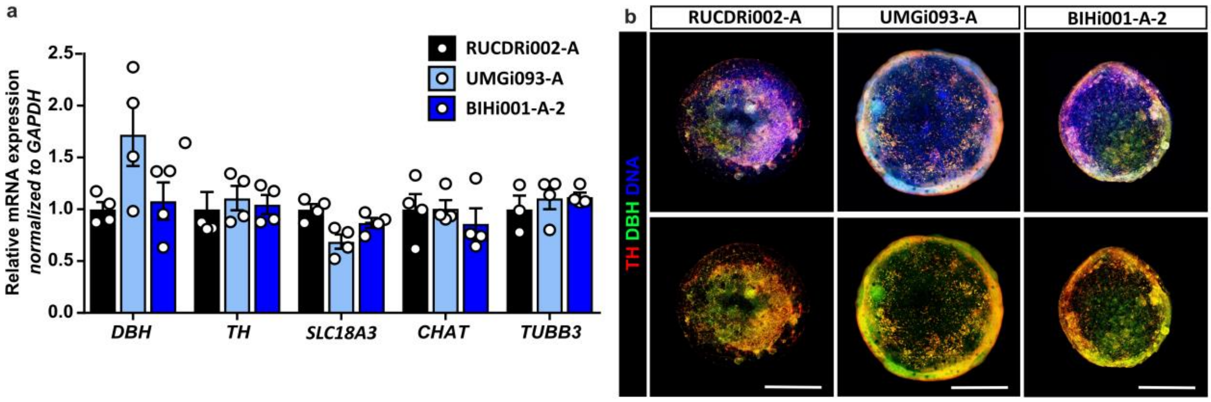


***Supplementary Figure 6| Reproducibility of SNO differentiation protocol in three iPSC-lines. a,*** *qPCR analysis of SN-marker DBH and TH, cholinergic marker SLC18A3 and CHAT as well as general neuron marker TUBB3 showed no difference between SNO from three different cell lines (RUCDRi002-A, BIHi001-A-2, UMGi093-A). Values were normalized to housekeeper and standard cell line RUCDRi002-A. b, WhIF of SNO from three different iPSC lines stained for DNA and SN-marker DBH and TH. Scale bar, 1 mm.*

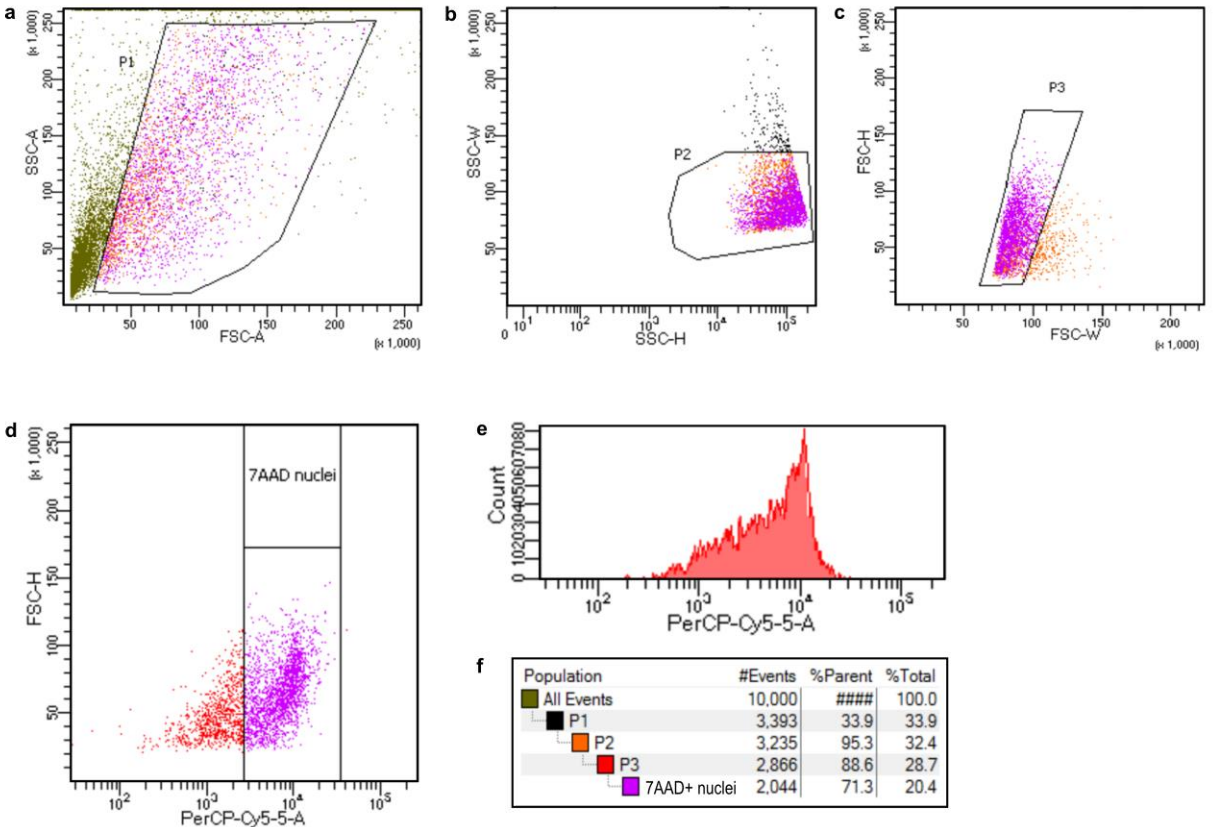


***Supplementary Figure 7| Gating strategy for sorting nuclei from frozen iEHM lysates using 7AAD staining. a-c,*** *Gating strategy for exclusion of debris (P1, P2, P3) and for 7AAD-stained nuclei* ***(d)*** *from iEHM lysates.* ***e,*** *Hierarchy of the gated populations including proportions.* ***f,*** *Histogram of raw nuclei count (PerCP-Cy5.5 signal, 7AAD dye).*

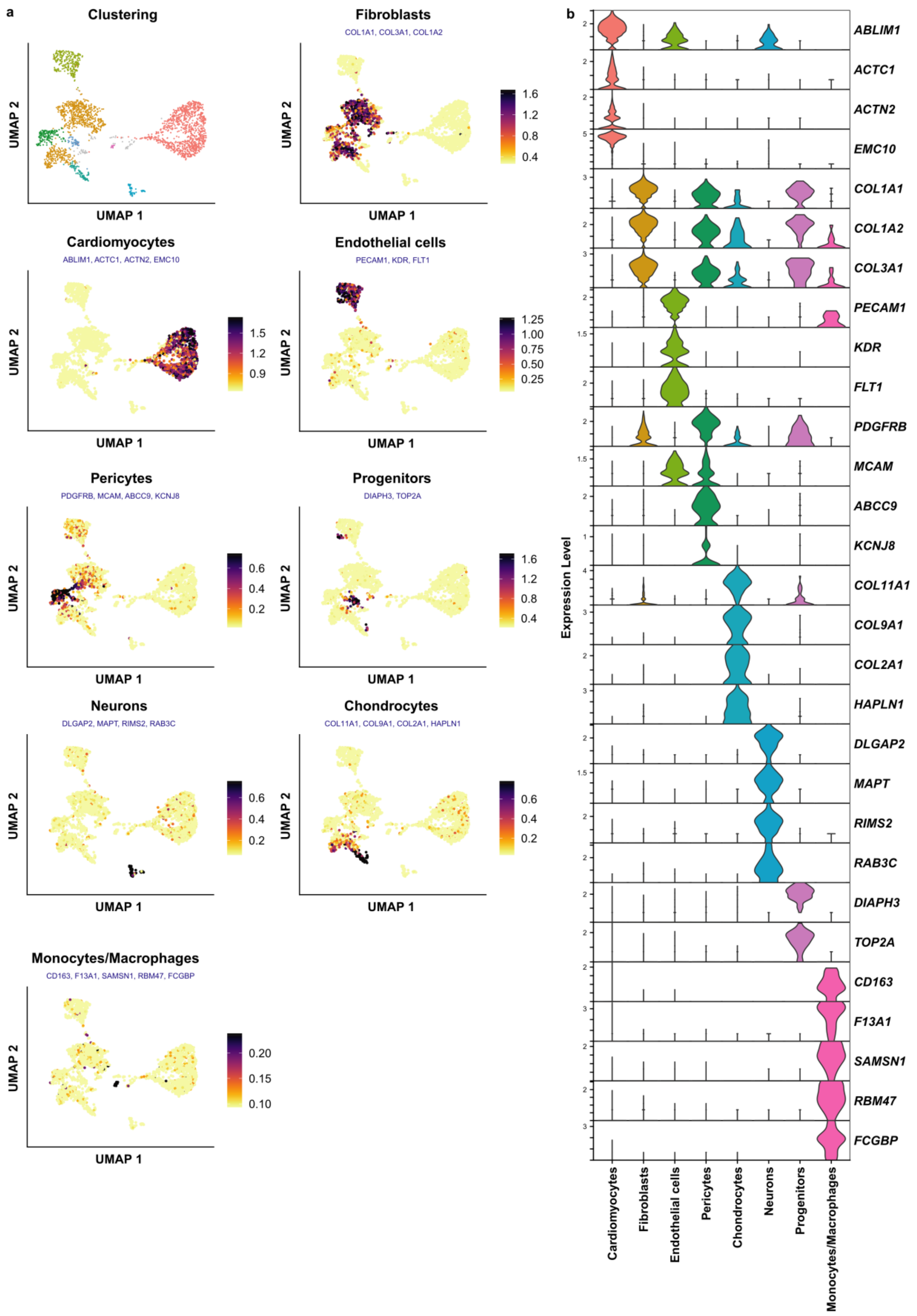


***Supplementary Figure 8| Cell cluster annotation of snRNA-sequencing. a,*** *UMAP plots depicting the consensus expression (average module score) of cell-type specific marker genes used to annotate the clusters in the dataset. Gene names are highlighted in blue under the plot title of the corresponding cell-type.* ***b,*** *Violin plots showing the normalized expression of individual marker genes from* ***a*** *across the annotated cell-types.*

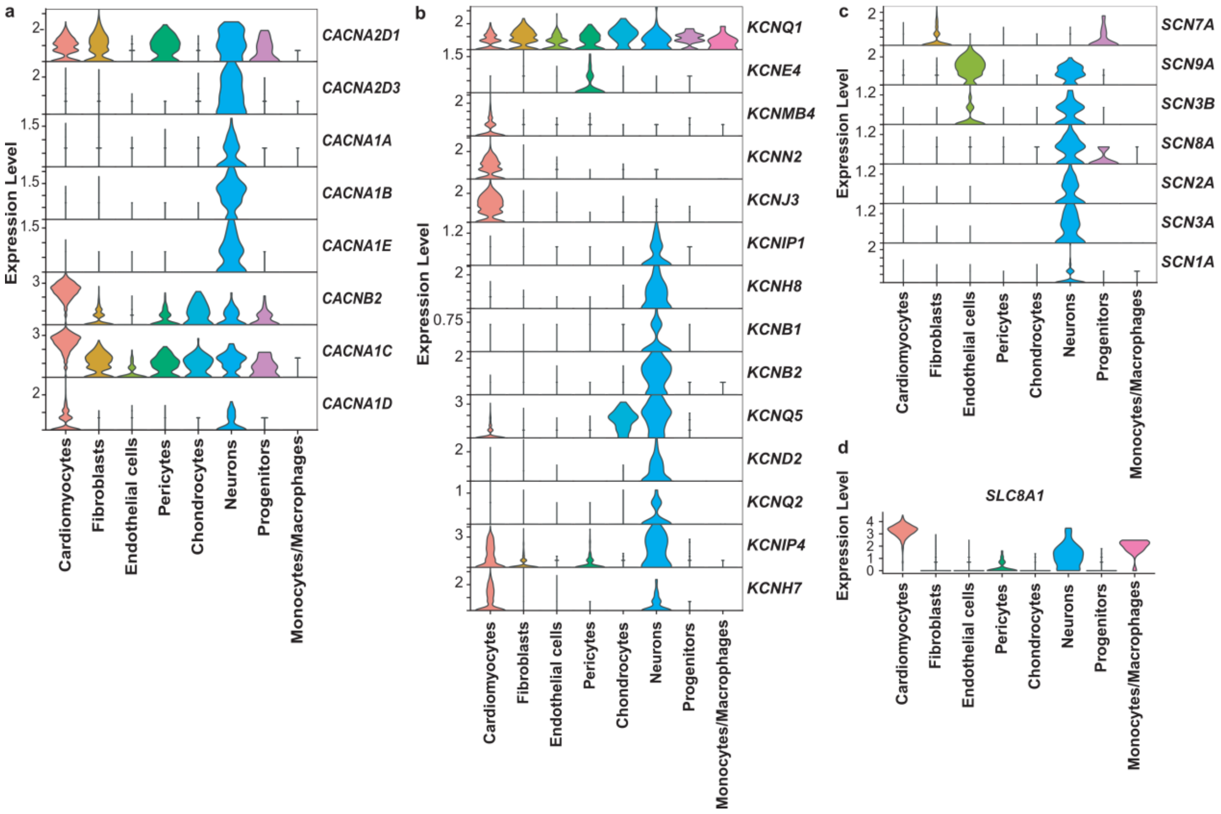


***Supplementary Figure 9| Expression analysis of ion channels across the cell types from snRNA-sequencing.*** *Violin plots showing the normalized expression of* ***a,*** *voltage-gated calcium (Ca$^{2+}$) channel transcripts,* ***b,*** *voltage-gated potassium (K$^{+}$) channel transcripts,* ***c,*** *voltage-gated sodium (Na$^{+}$) channel transcripts and* ***d,*** *the Na+/Ca2+ exchanger SLC8A1 transcript, across the annotated cell-types.*

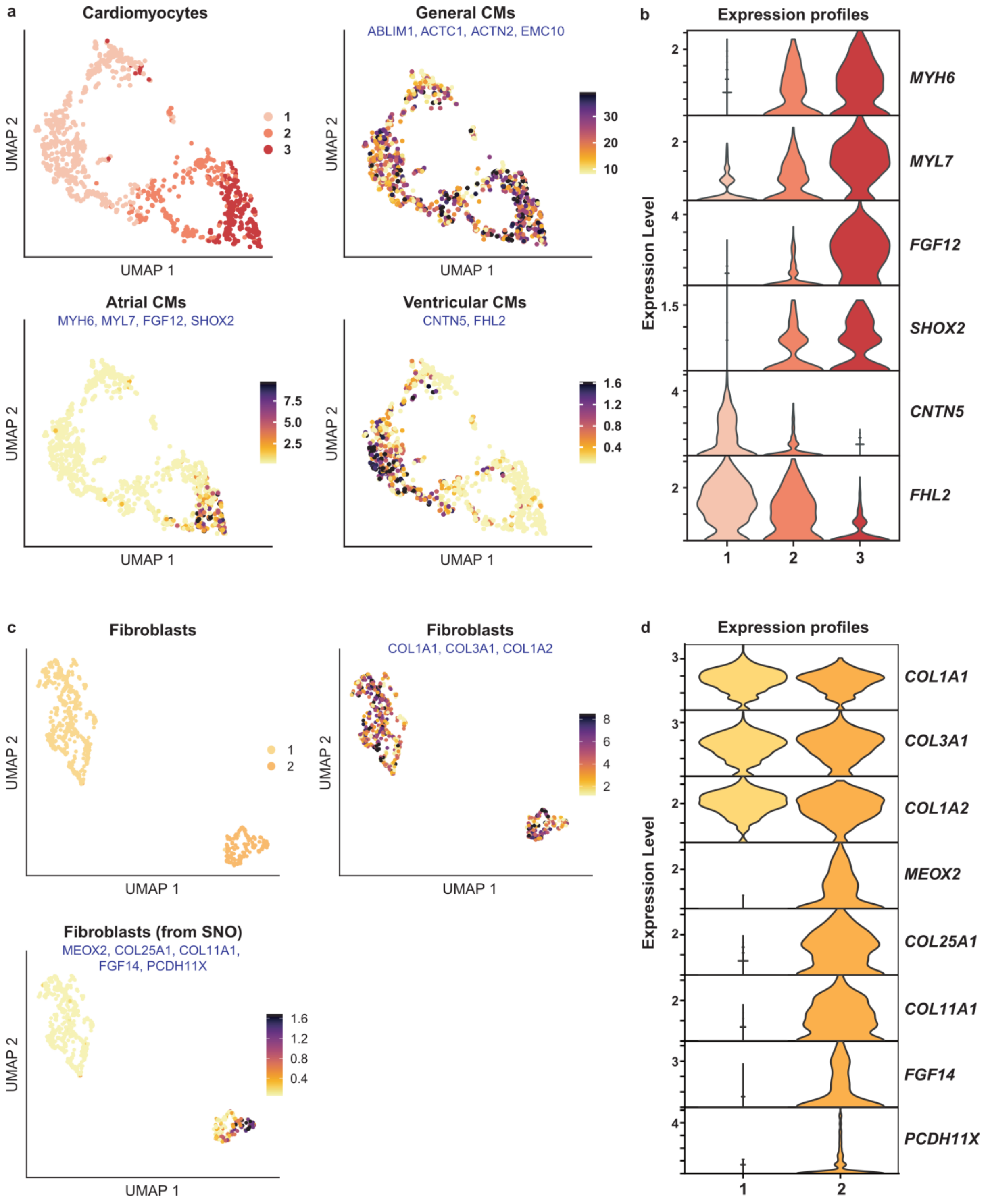


***Supplementary Figure 10| Analysis of cardiomyocyte and fibroblast subcluster in the iEHM. a,*** *UMAP plots depicting the consensus expression (average module score) of atrial, ventricular and general cardiomyocyte marker genes across the cardiomyocyte sub-clusters (UMAP plot in the top-left panel). Gene names are highlighted in blue under the plot title of the corresponding sub-type.* ***b,*** *Violin plots showing the normalized expression of individual atrial and ventricular cardiomyocyte marker genes from* ***a*** *across the three sub-clusters.* ***c,*** *UMAP plots depicting the consensus expression (average module score) of fibroblast marker genes across the fibroblast sub-clusters (UMAP plot in the top-left panel).* ***d,*** *Violin plots showing the normalized expression of individual fibroblast marker genes from* ***c*** *across the two sub-clusters.*

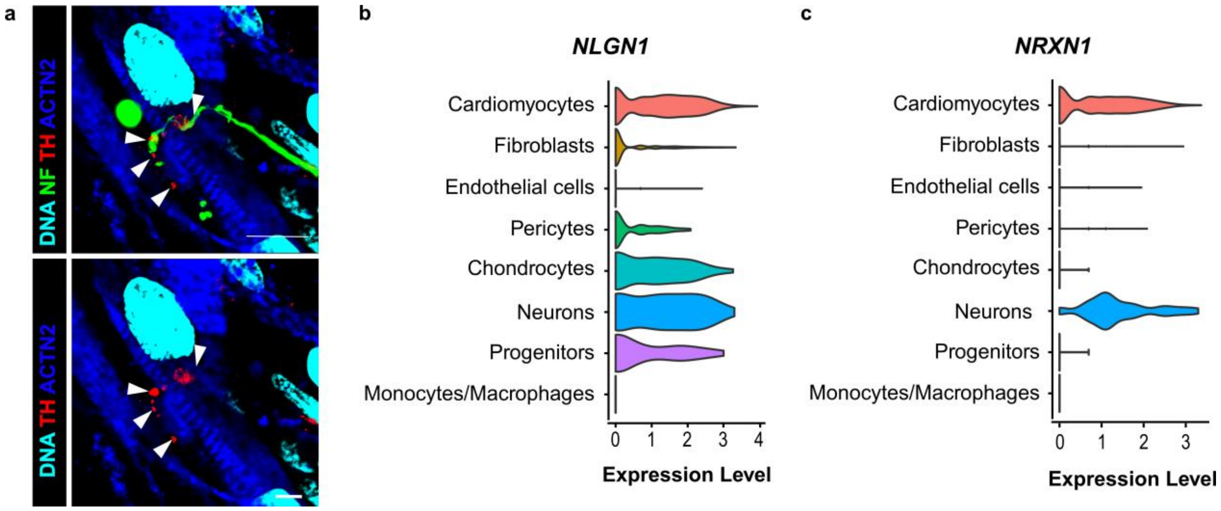


***Supplementary Figure 11| Investigation of neuro-cardiac junctions in iEHM. a**, WhIF showing co-staining of synaptic bouton-like varicosity (NF, green) co-localizing with autonomic neuron marker TH (red) on the surface of cardiomyocytes (ACTN2, blue). Co-localization is indicated by white arrowheads. Scale bar, 10 µm. **b**, Expression of postsynaptic adhesion molecule neuroligin 1 (NLGN1) across different cell types in the iEHM. **c**, snRNA sequencing expression analysis of presynaptic marker neurexin 1 (NRXN1) in the annotated iEHM cell clusters (see **Supplementary Figure 8**).*

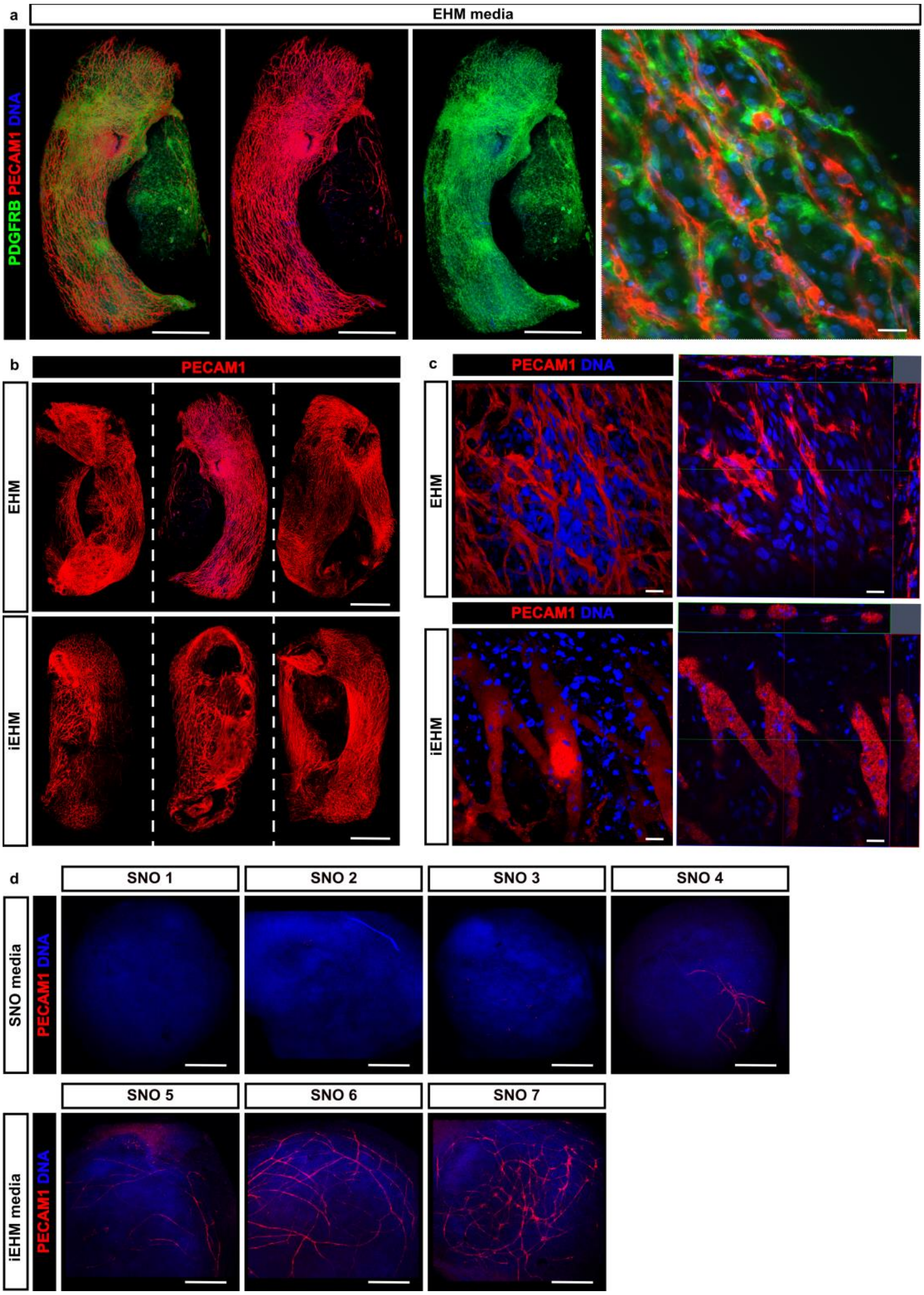


***Supplementary Figure 12| Vascular network development in SNO, EHM and iEHM. a****, WhIF of 6 weeks EHM cultured in EHM medium reveals a dense endothelial network (PECAM1, red) and presence of PDGFRb (green)-positive pericytes. Scale bar, 1 mm.* ***(Right)*** *Close-up view showing close localization of pericytes and endothelial cells. Scale bar, 20 µm.* ***b****, PECAM1 (red) immunofluorescence analysis of three representative EHM and iEHM shows a similar extent of vascularisation of 6 week old EHM and iEHM. Scale bar, 1 mm.* ***c****, Close-up view of PECAM1 staining in EHM and iEHM* ***(left)*** *and the 3D-reconstruction* ***(right)*** *show a larger vessel diameter in some areas of the iEHM when compared to EHM. Scale bar, 20 µm.* ***d, (Top)*** *WhIF of d93 SNO cultured exclusively in SNO media shows presence of endothelial cells in only one out of four tissues.* ***(Bottom)*** *After eight weeks of culture in iEHM medium, d103 SNO show the presence of endothelial cells in all of the screened tissues. Scale bar, 200 µm.*

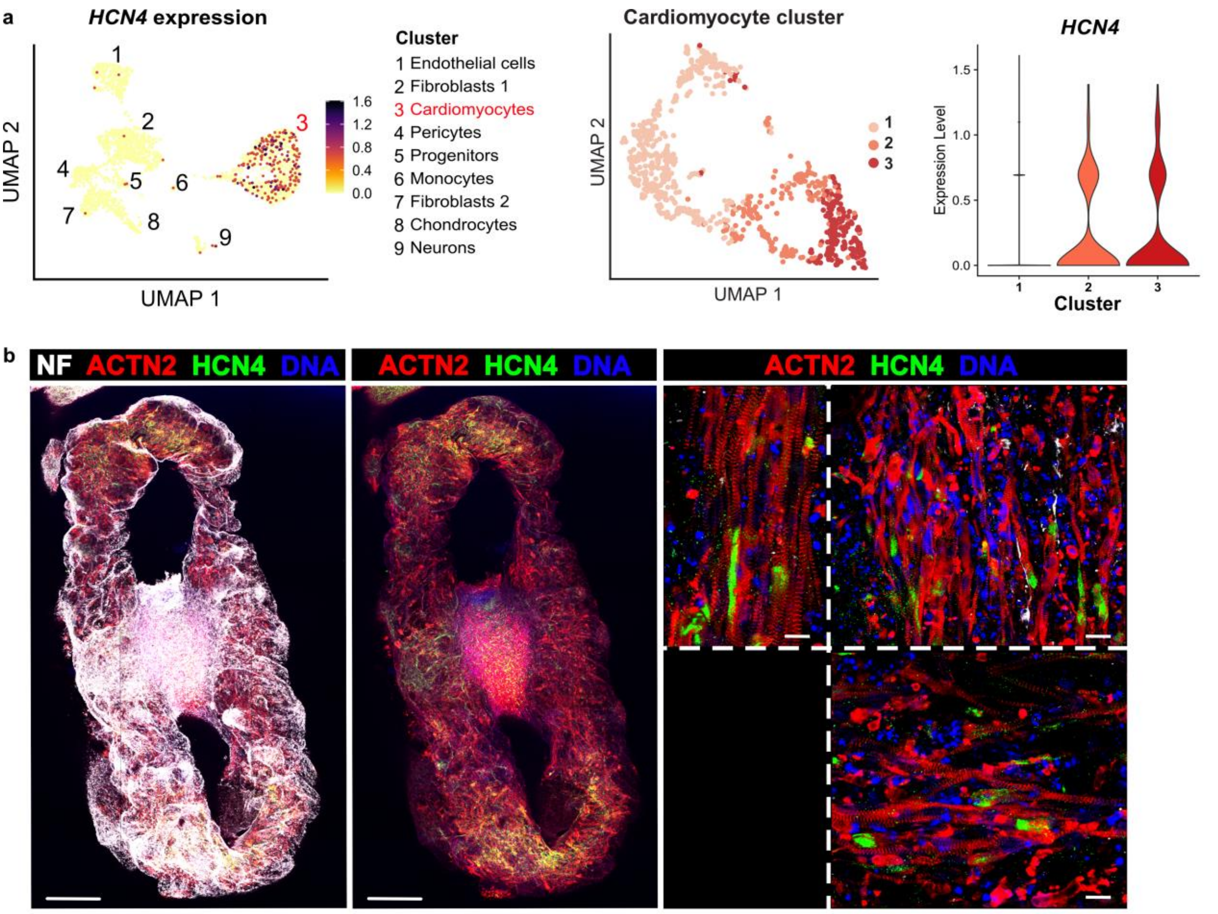


***Supplementary Figure 13| Identification of pacemaker-like cells in iEHM a**, **(Left)** UMAP plots showing the distribution of cells expressing the pacemaker-like-cell marker HCN4 in the clusters in the dataset including cluster annotations (compare **Supplementary Figure 8**). **(Right)** Expression of HCN4 in the three cardiomyocyte cluster (Supplementary Figure 10)**. b**, WhIF of iEHM stained for NF (grey), cardiomyocyte marker ACTN2 (red), pacemaker-like-cell marker HCN4 (green) and DNA (blue). **(Left)** Overview images show pacemaker-like-cells dispersed throughout the innervated areas of the EHM. Scale bar, 500 µm. **(Right)** Close-up view show HCN4 positive cells in between ACTN2-positive CM co-localizing with NF-positive neurites (top left panel). Scale bar, 20 µm.*

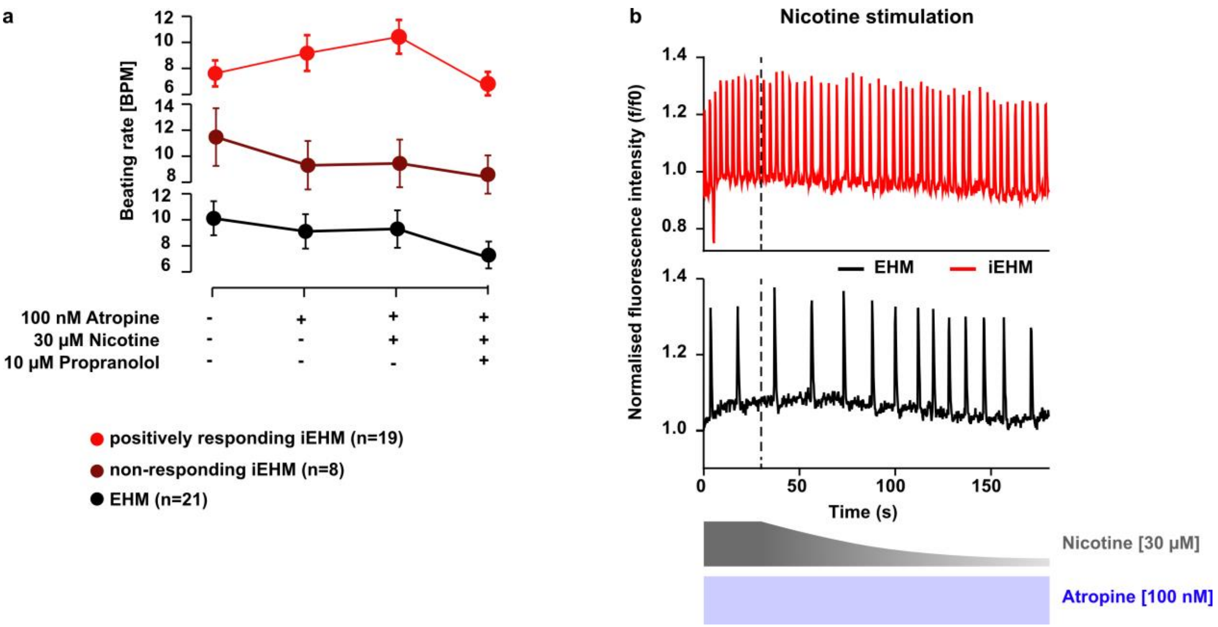


***Supplementary Figure 14| Details of pharmacological stimulation of iEHM. a**, Chronotropic response to stimulation of EHM and iEHM with 100 nM atropine, 30 µM nicotine and 10 µM propranolol. iEHM could be separated into two distinct groups of responding (N=19) and non-responding tissues (N=8) that behaved similar to EHM controls. **b**, Example traces showing the acute effect of nicotine stimulation on EHM and iEHM mean fluorescence intensity (f/f0) of the ROI (see in **Figure 3e**) in the presence of 100 nM atropine.*

### Supplementary Material

***Supplementary Table 4.*** *Detection parameters for a second compound-specific mass transition used as qualifier. Q1=first quadrupole, Q3=third quadrupole, DP=declustering potential, CE=collision energy, CXP=collision cell exit potential.*

| Compound | Mass Q1 [Da] | Mass Q3 [Da] | DP [V] | CE [V] | CXP [V] |
|---|---|---|---|---|---|
| Noradrenaline | 170.2 | 152.0<br>(107.0) | 36 | 10<br>-28 | 10<br>-20 |
| Dopamine | 154.1 | 137.2<br>(91.0) | 36 | 15<br>-33 | 10<br>-8 |
| Acetylcholine | 147.0 | 87.0<br>(88.0) | 36 | 19<br>-20 | 16<br>-16 |
| Noradrenaline-d6 | 176.1 | 158.0<br>(111.0) | 31 | 11<br>-29 | 10<br>-6 |
| Buformin | 157.91 | 60.9<br>(47.0) | 36 | 35<br>-66 | 10<br>-8 |
| Choline-d9 | 113.108 | 69.1<br>(66.1) | 66 | 27<br>-44 | 12<br>-12 |

***Supplementary Table 5.*** *List of primer for quantitative real-time PCR.*

| Gene | Forward primer | Reverse primer | Amplicon length (bp) |
|---|---|---|---|
| *ASCL1* | CGGTCTCATCCTACTCGTCG | GTTGTGCGATCACCCTGCTT | 150 |
| *CHAT* | GGATCGCTGGACATGATTG | CCACACCGCCAGGTCGC | 103 |
| *DBH* | TGGGTGCCAAGGCATTTTAC | ATGCCTGAGGAGTCGTTTCG | 136 |
| *GAD1* | AGGCAATCCTCCAAGAACC | TGAAAGTCCAGCACCTTGG | 218 |
| *GAPDH* | CCTCAAGATCATCAGCAATGCC | ATGTTCTGGAGAGCCCCGC | 189 |
| *GATA2* | GCAGAACCGACCACTCATCA | AGTGGCCTGTTAACATTGTGC | 177 |
| *GBX2* | GTTCCCGCCGTCGCTGATGAT | GCCGGTGTAGACGAAATGGCCG | 118 |
| *GLAST* | CCCCTTACAAAATCAGAAAAGTTGT | CCCATCTTGGGCTCTTCTCC | 151 |
| *MNX1* | TGCCTAAGATGCCGACTTC | AATCTTCACCTGGGTCTCGG | 194 |
| *OLIG2* | GCTGCGTCTCAAGATC | AGTCGCTTCATCTCCT | 192 |
| *PAX6* | CCCCACATATGCAGACACAC | TCACTTCCGGGAACTTGAAC | 112 |
| *PHOX2B* | CGCCGCAGTTCCTTACAAAC | TGTCGGGGTAGTGAGTCTCC | 140 |
| *PRPH* | TAAATATAAAGACGACTGTGCCTGA | GCAGAAGACTTGTCCAGCTCA | 151 |
| *SLC17A6* | GTAGACTGGCAACCACCTCC | CCATTCCAAAGCTTCCGTAGAC | 129 |
| *SLC18A2* | TTCCGACTGTCCCAGTGAAG | TGCTGGAGAAGGCAAACATAAT | 200 |
| *SLC6A2* | CAGGTTCAGCAACGACATCCAG | GTCGTAGGTGAGTGGCTTGAAG | 142 |
| *TH* | CGGGGCTTCTCGGACCAGGTGTA | CTCCTCGGCGGTGTACTCCACA | 111 |
| *TUBB3* | CAGCGTCTACTACAACGAGG | AGGCCTGAAGAGATGTCCAA | 121 |
| *SLC18A3* | CTGCTAGTGAACCCCTTGAGC | CAGGACTGTAGAGGCGAACAT | 99 |

***Supplementary Table 6**. List of primary and secondary antibodies.*

| Antigen | Modification | Brand | Cat. No. | Host | Dilution | Type |
|---|---|---|---|---|---|---|
| ACTN2 | None | Sigma | A7811 | Mouse | 1:1000 | Primary |
| BRN3A | None | Merck Millipore | MAB1585 | Mouse | 1:500 | Primary |
| CTNT | None | Abcam | ab45932 | Rabbit | 1:250 | Primary |
| DBH | None | Sigma | HPA070789 | Rabbit | 1:100 | Primary |
| HCN4 | None | Abcam | ab69054 | Rabbit | 1:200 | Primary |
| Neurexin 1 | None | Sigma | HPA071400 | Rabbit | 1:200 | Primary |
| NF (H) | None | Synaptic Systems | 171 106 | Chicken | 1:100 | Primary |
| NF (H) | None | Biolegend | 822601 | Chicken | 1:1000 | Primary |
| PAX6 | None | Biolegend | 901301 | Mouse | 1:500 | Primary |
| PDGFRb | None | Cell Signalling | 28E1 | Rabbit | 1:100 | Primary |
| PECAM1 | None | DAKO | M0823 | Mouse | 1:100 | Primary |
| Peripherin | None | NOVUS | NBP1-05423 | Chicken | 1:1000 | Primary |
| Phalloidin | Alexa Fluor 488 | Invitrogen | A12379 | - | 1:200 | Coupled |
| PHOX2B | None | Santa Cruz | sc-376997 | Mouse | 1:250 | Primary |
| Synapsin 1 | None | Synaptic systems | 106 011 | Mouse | 1:200 | Primary |
| TH | None | Synaptic Systems | 213 004 | Guinea pig | 1:500 | Primary |
| vAChT | None | Synaptic Systems | 139 103 | Rabbit | 1:500 | Primary |
| VGLUT1 | Abberior STAR 635P | NanoTag | N1602-Abb635P | Rabbit | 1:250 | Coupled |
| Mouse | Alexa Fluor 633 | Invitrogen | A21052 | Goat | 1:400 | Secondary |
| Chicken | Alexa Fluor 488 | Invitrogen | A11039 | Goat | 1:400 | Secondary |
| Rabbit | Alexa Fluor 546 | Invitrogen | A11035 | Goat | 1:400 | Secondary |
| Guinea Pig | Alexa Fluor 546 | Invitrogen | A11074 | Goat | 1:400 | Secondary |

### Supplementary Videos

**Supplementary Video 1.** Spontaneous beating of iEHM 1 week after fusion

**Supplementary Video 2.** Representative immunofluorescence picture depicting neurofilament positive axons (green) interlacing alpha-actinin positive cardiomyocytes (red) in iEHM

**Supplementary Video 3.** Representative immunofluorescence picture 1 depicting synapsin positive pre-synaptic terminals (red) of noradrenergic neurons (TH, blue) in between cardiomyocytes (cTNT, green) in 8 week-old iEHM.

**Supplementary Video 4.** Representative immunofluorescence picture 2 depicting synapsin positive pre-synaptic terminals (red) of noradrenergic neurons (TH, blue) in between cardiomyocytes (cTNT, green) ) in 8 week-old iEHM.

**Supplementary Video 5.** Representative immunofluorescence picture 3 depicting synapsin positive pre-synaptic terminals (red) of noradrenergic neurons (TH, blue) in between cardiomyocytes (cTNT, green) ) in 8 week-old iEHM.

**Supplementary Video 6.** Representative immunofluorescence picture 4 depicting synapsin positive pre-synaptic terminals (red) of noradrenergic neurons (TH, blue) in between cardiomyocytes (cTNT, green) ) in 8 week-old iEHM.

**Supplementary Video 7.** Representative immunofluorescence picture 5 depicting synapsin positive pre-synaptic terminals (red) of noradrenergic neurons (TH, blue) in between cardiomyocytes (cTNT, green) in 8 week-old iEHM.

**Supplementary Video 8.** Representative immunofluorescence picture 6 depicting synapsin positive pre-synaptic terminals (red) of noradrenergic neurons (TH, blue) in between cardiomyocytes (cTNT, green) in 8 week-old iEHM.

**Supplementary Video 9.** Representative immunofluorescence picture 1 depicting PECAM1 positive vascular network (red) surrounded by PDGFRb positive pericytes (cyan) in close proximity to neurons (NF, green) in 8 week-old iEHM.

**Supplementary Video 10.** Representative immunofluorescence picture 2 depicting PECAM1 positive vascular network (red) surrounded by PDGFRb positive pericytes (cyan) in close proximity to neurons (NF, green) in 8 week-old iEHM.

**Supplementary Video 11.** Representative immunofluorescence picture 3 depicting PECAM1 positive vascular network (red) surrounded by PDGFRb positive pericytes (cyan) in close proximity to neurons (NF, green) in 8 week-old iEHM.

**Supplementary Video 12.** Chronotropic response of EHM and optogenetic iEHM evoked by light stimulation
